# Age-dependent NMDA receptor function is regulated by the Amyloid Precursor Protein

**DOI:** 10.1101/2022.07.20.500736

**Authors:** Joana Rajão-Saraiva, Mariana Temido-Ferreira, Joana E. Coelho, Aurore Ribera, Sébastien Moreno, Michael Willem, Hélène Marie, Luísa V. Lopes, Paula A. Pousinha

## Abstract

N-methyl-D-aspartate receptors (NMDARs) are critical for the maturation and plasticity of glutamatergic synapses. In the hippocampus, NMDARs mainly contain GluN2A and/or GluN2B regulatory subunits. The amyloid precursor protein (APP) has emerged as a putative regulator of NMDARs, but the impact of this interaction to their function is largely unknown.

By combining patch-clamp electrophysiology and molecular approaches, we unravel a dual mechanism by which APP controls GluN2B-NMDARs, depending on the life stage. We show that APP is highly abundant specifically at the postnatal postsynapse. It interacts with GluN2B-NMDARs, controlling its synaptic content and mediated currents, both in infant mice and primary neuronal cultures. Upon aging, the APP amyloidogenic derived C-terminal fragments, rather than APP full length, contribute to aberrant GluN2B-NMDAR currents. Accordingly, we found that the APP processing is increased upon aging, both in mice and human brain, and interfering with this mechanism normalized the aberrant currents.

While the first mechanism might be essential for synaptic maturation during development, the latter could contribute to age-related synaptic impairments.

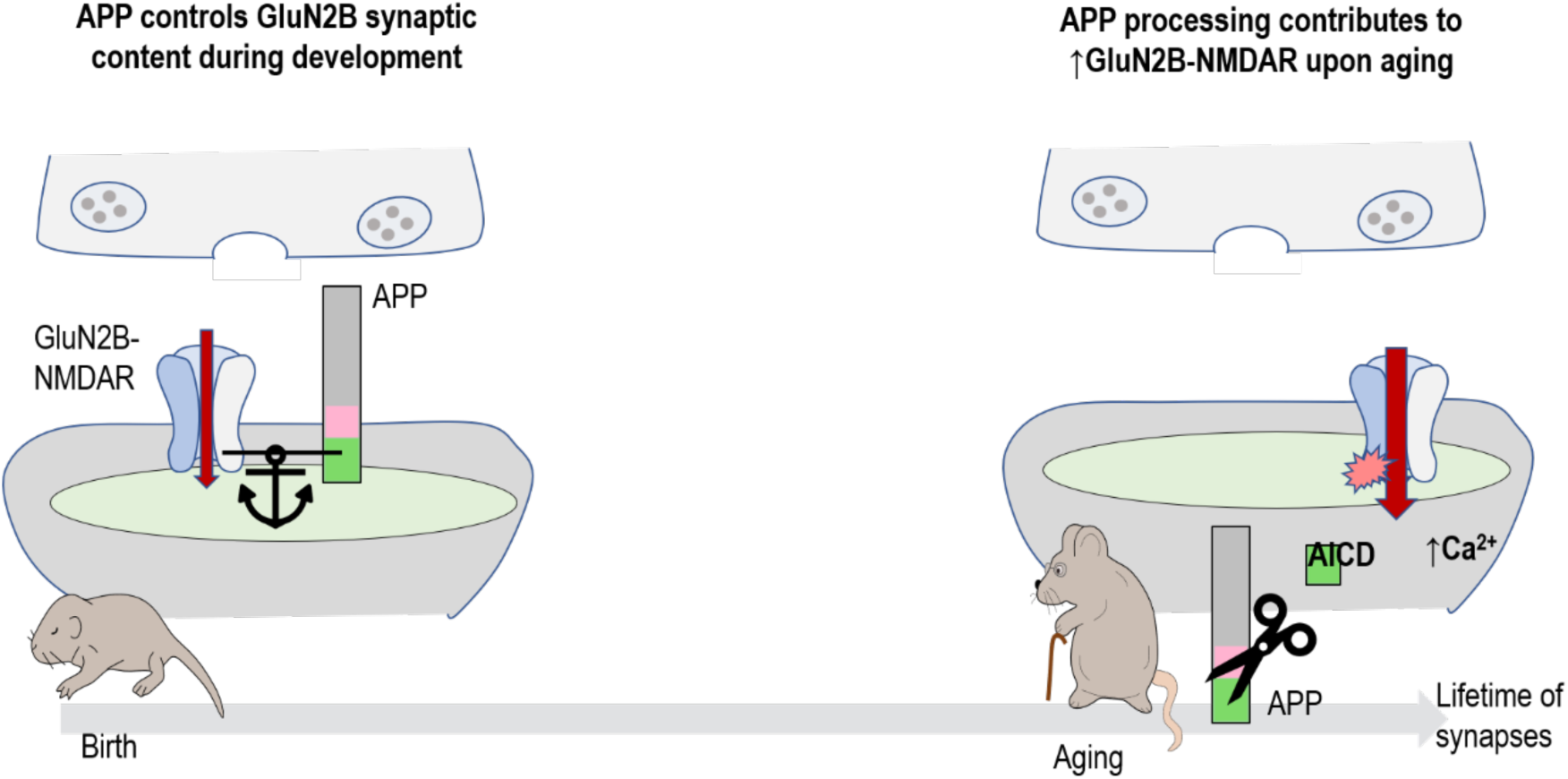

**In brief:** Rajão-Saraiva et al. identified the APP full-length protein as a regulator of glutamatergic transmission in immature synapses, by controlling GluN2B synaptic content and mediated currents during postnatal development. Upon aging, the APP amyloidogenic derived C-terminal fragments, rather than APP full length, contribute to aberrant GluN2B-NMDAR currents. Our work highlights the importance of keeping APP processing under tight control, to ensure the normal functioning of glutamatergic synapses, being particularly relevant to understand age-related synaptic impairments and AD.

## Introduction

The activation of N-methyl-D-aspartate receptors (NMDARs) at glutamatergic synapses results in calcium influx into neurons, activating downstream signalling pathways (Forsythe & Westbrook, 1988, Hardingham et al., 1999). Due to their voltage-dependent block by magnesium, NMDARs act as coincidence detectors for pre and postsynaptic activity since receptor activation requires glutamate release and strong membrane depolarization (Nowak et al., 1984). This process is critical for the formation, maturation, maintenance, and plasticity of glutamatergic synapses, thereby contributing to development, learning and memory processes (Gambrill & Barria, 2011, Morris et al., 1986). On the other hand, NMDAR dysfunction is associated with several pathological conditions, including neurodevelopmental disorders (Burnashev & Szepetowski, 2015) and aged-related neurodegenerative diseases such as Alzheimer’s Disease (AD) (Zhou & Sheng, 2013, Parameshwaran et al., 2008).

One of the main factors that determines NMDARs properties is their subunit composition. NMDARs are heterotetramers composed of two obligatory GluN1 subunits and two GluN2 (A–D) or GluN3 (A and B) subunits (Paoletti et al., 2013). In the hippocampus and cortex, NMDARs mainly contain GluN2A and/or GluN2B regulatory subunits (Monyer et al., 1994, Al-Hallaq et al., 2007, Gray et al., 2011). GluN2B-NMDARs differ from the GluN2A subtype due to their slower deactivation kinetics, higher Ca^2+^ charge per unit of current and specific intracellular interactors (Sobczyk et al., 2005, Vicini et al., 1998, Erreger et al., 2005, Strack & Colbran, 1998). Therefore, the synaptic GluN2B/GluN2A ratio determines the consequences of NMDAR activation, including total calcium influx and downstream signalling. This ratio is not static, changing in response to neuronal activity and sensory experience during postnatal development, but also in adult synapses (Bellone & Nicoll, 2007, He et al., 2006, Quinlan et al., 1999, Philpot et al., 2001, Yashiro et al., 2005, Baez et al., 2013).

During development, glutamatergic synaptic transmission in nascent synapses is mainly mediated by GluN2B-NMDARs (Durand et al., 1996, Hsia et al., 1998, Bellone & Nicoll, 2007, Flint et al., 1997). When there is strong or synchronous neuronal activity, resulting in enough calcium entry through GluN2B-NMDARs (Durand et al., 1996, Gray et al., 2011, Adesnik et al., 2008), synapses undergo maturation, leading to alterations in the composition of postsynaptic receptors, namely a shift from a predominance of GluN2B to GluN2A-NMDARs (Williams et al., 1993, Bar-Shira et al., 2015, Law et al., 2003, Jantzie et al., 2015). Upon adulthood, both NMDAR subtypes contribute for normal calcium signalling (Sobczyk et al., 2005), synaptic plasticity (Massey et al., 2004, Barria & Malinow, 2005) (Yashiro & Philpot, 2008, Philpot et al., 2007) and memory formation (von Engelhardt et al., 2008, Bannerman et al., 2008). However, these NMDAR-dependent processes become dysregulated upon aging, resulting in elevated postsynaptic calcium levels (Thibault et al., 2001), alterations in long-term potentiation/depression (LTP/LTD) (Burke & Barnes, 2006, Temido-Ferreira et al., 2020) and memory deficits (Tombaugh et al., 2002). Based on this, NMDAR gain of function has emerged as a possible explanation for age-related synaptic impairments (Kumar, 2015). Accordingly, GluN2B present higher retention at aged synapses (Zamzow et al., 2013, al Abed et al., 2020) and an inverse correlation with memory performance (Zhao et al., 2009). Moreover, GluN2B-NMDAR overactivation and consequent excessive calcium influx are known to contribute to age-associated neurodegenerative disorders (D. G. Ferreira et al., 2017, Hanson et al., 2015, Li et al., 2011). This hypothesis is counterintuitive, as the GluN2B subunit is thought to be the most affected by aging (Magnusson, 2012), showing a decline in expression levels (Magnusson, 2000, Zhao et al., 2009, Clayton & Browning, 2001, Sonntag et al., 2000, Bai et al., 2004). Further knowledge by electrophysiological examination of functional changes in GluN2B-NMDARs during the aging process may be the key to understand aging cognitive disabilities.

The amyloid precursor protein (APP) has emerged as a putative NMDAR regulator, since it interacts with NMDARs in rodent brain lysates and in primary neuronal cultures (Cousins et al., 2009, Hoe et al., 2009). Also, we previously showed that in utero silencing of APP causes the loss of GluN2B synaptic contribution in infant mice (Pousinha et al., 2017). Studying the role of APP is challenging considering the functional redundancy with members of the same protein family, but also given the multiple APP fragments, generated by secretase cleavage, with specific cellular functions (Müller et al., 2017). The APP family is composed of three highly conserved transmembrane glycoproteins, the APP and amyloid precursor-like proteins 1 and 2 (APLP1, APLP2), with overlapping functions (Müller & Zheng, 2012). APP undergoes extracellular cleavage mainly by α- or β-secretase (non-amyloidogenic or amyloidogenic pathway, respectively), resulting in the formation of large N-terminal extracellular fragments of secreted APP and smaller membrane-bound C-terminal fragments (Müller et al., 2017). Subsequently, the C-terminal fragments are subjected to an intramembranous scission by the γ-secretase complex (Wolfe et al., 1999) to generate the APP intracellular domain (AICD) and to simultaneously release p3 (after α-secretase cleavage) or the AD related amyloid-β peptide (Aβ, after β-secretase cleavage). Therefore, APP has been mostly studied in the context of AD, while far less is known about its physiological role throughout life. APP levels reach their peak during brain development (Kirazov et al., 2001, Löffler & Huber, 1992), when it has been suggested to participate in synaptogenesis and clustering of synaptic proteins, as revealed by *in vitro* overexpression/knockdown studies (Wang et al., 2009, K. J. Lee et al., 2010). Later in life, a role for APP in synaptic plasticity has been proposed, considering the impairments in LTP and cognitive behavioural performance observed in aged APP-knockout animals (>10months), but not in earlier stages (2-4 months) (Dawson et al., 1999, Tyan et al., 2012).

We now explored the mechanism by which APP regulates NMDAR synaptic transmission at different life stages. Combining electrophysiological outputs and molecular approaches, we found a dual mechanism by which APP controls GluN2B-NMDARs. We identified the APP full-length protein as a new regulator of glutamatergic transmission in immature synapses, by controlling GluN2B synaptic content and mediated currents during development. In addition, we gathered strong evidence showing that APP C-terminal fragments generated by the amyloidogenic pathway modify NMDAR transmission, favouring the GluN2B synaptic contribution at later life stages. Our work highlights the importance of keeping APP processing under tight control, to ensure the normal functioning of glutamatergic synapses, being particularly relevant to understand age-related synaptic impairments and AD.

## Results

### GluN2B-NMDAR synaptic contribution is increased in infant and aged mice

We recorded pharmacologically isolated NMDAR excitatory post-synaptic currents (EPSCs) in CA1 pyramidal cells in the hippocampus, evoked by electrical stimulation of Schaffer collaterals (**Figure 1a**) in C57BL/6 mice at different ages: infant (7-10 days), adult (10 – 16 weeks) and aged (18 – 20 months). To investigate if age-mediated alterations of NMDAR currents were due to modifications of NMDAR subunit composition, we measured the time constant for the weighted component (τ_weighted_) of NMDAR EPSC deactivation kinetics. We found that NMDAR EPSCs deactivation kinetics are age-dependent, since the τ_weighted_ was higher in infant (156,2 ms ± 4,20, mean ± SEM) and aged mice (177,0 ms ± 12,66), compared to the reference group of adults (126,3 ms ± 9,11) (**Figure 1b, c**). We also found an increase in the relative amplitude of the slow component (A_slow_) for infant and aged mice (**Supplementary Figure 1a**).

**Figure 1 -.**
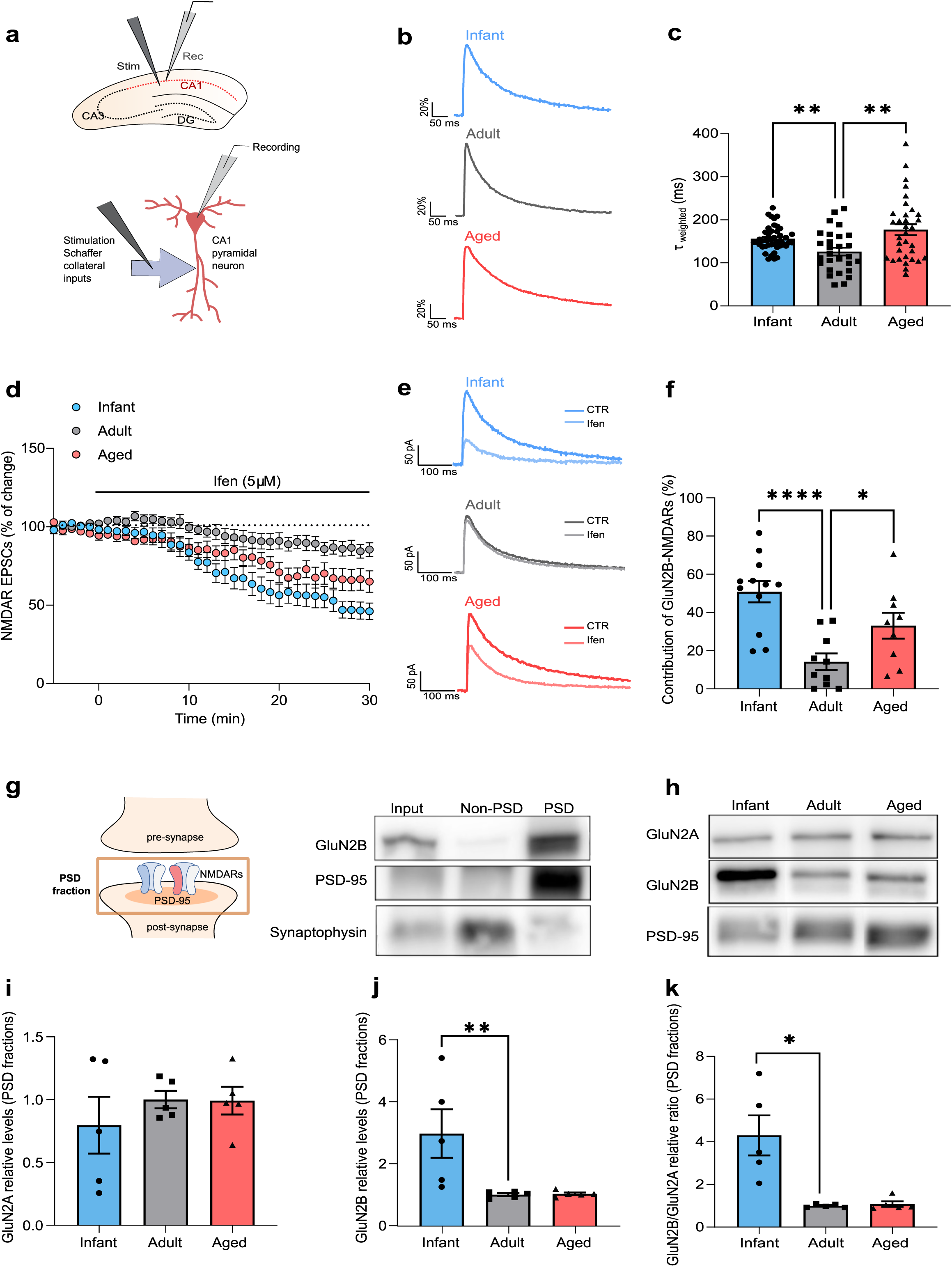
GluN2B-NMDAR synaptic contribution is increased in infant and aged mice. (a) Schematic diagram showing the locations of stimulating and recording electrodes in the hippocampus and in a CA1 pyramidal neuron for whole cell patch-clamp experiments. (b) Comparison of representative whole-cell patch-clamp recordings of pharmacologically isolated NMDAR EPSCs, normalized to the peak amplitude (in %), from infant, adult and aged mice, illustrating differences in deactivation kinetics. (c) The weighted time constant (τ_weighted_) was calculated using the relative contribution of both slow/fast components of NMDAR EPSCs and reflects the overall deactivation kinetics. Results are expressed as the mean ± SEM (Kruskal Wallis followed by an Uncorrected Dunn’s multiple comparisons test using the adult group as reference, **p<0.01, n=28-44). (d) Time course of ifenprodil (5µM) effect on pharmacologically isolated NMDAR EPSC amplitude in CA1 pyramidal neurons, measured by whole-cell patch clamp in infant, adult and aged C57BL/6 wild-type mice. Results are expressed as the mean ± SEM (n=9-12). (e) Traces show NMDAR EPSCs recorded before (CTR) and after 30 min of Ifenprodil 5 µM perfusion (Ifen). (f) GluN2B contribution was calculated as the percentage of change in NMDAR EPSC amplitude after 30 min of ifenprodil perfusion. Results are expressed as the mean ± SEM (One-way ANOVA followed by an Uncorrected Fisher’s LSD’s multiple comparisons test using the adult group as reference, *p<0.05, ****p<0.0001, n=9-12). (g) Schematic representation and immunoblotting analysis of PSD-enriched fractions from the hippocampal tissue of adult C57BL/6 wild-type mice. Membranes were immunoblotted with antibodies for GluN2B, PSD-95 and synaptophysin. PSD-fractions are enriched in PSD-95, whereas non-PSD fractions contain high levels of synaptophysin. (h) Representative western blot of hippocampal lysates subjected to biochemical fractionation to obtain PSD-enriched fractions from infant, adult and aged C57BL/6 mice. Membranes were immunoblotted with antibodies for GluN2A, GluN2B and PSD-95. (i, j) Results from PSD-enriched fractions were normalized with PSD-95 and are expressed as the mean ± SEM (One-way ANOVA followed by an Uncorrected Fisher’s LSD’s multiple comparisons test using the adult group as reference, **p<0.01, n=5). (k) Results show the relative GluN2B/GluN2A in PSD-enriched fractions and are expressed as the mean ± SEM (Kruskal Wallis followed by an Uncorrected Dunn’s multiple comparisons test using the adult group as reference, *p<0.05, n=5).

Since GluN1/GluN2B heterodimers display slower deactivation kinetics than GluN1/GluN2A heteromers (Vicini et al., 1998), this suggests that the GluN2B contribution to NMDAR EPSCs is higher at infant and aged life stages, when compared to adults. To test this hypothesis, we evaluated the effect of the selective GluN2B inhibitor, ifenprodil (5µM), on NMDAR EPSCs. We found that the contribution of GluN2B to NMDAR EPSCs is of 50,88% ± 5,58 in infants, decreased to 14,20% ± 4,32 in adults and increased to 33,11% ± 6,72 (mean ± SEM) in aged mice (n=9-12), as depicted in **Figure 1 d-f**.

We then correlated these effects with the levels of NMDAR subunits in fractions enriched in the postsynaptic density (PSD-enriched fractions) (**Figure 1g**). GluN2B levels exhibit a peak in infant mice and a decline after development, whereas the GluN2A profile is more stable at all ages (n=5) (**Figure 1h-k**). This leads to a synaptic ratio of GluN2B/GluN2A approximately 4 times higher in infant mice, than in adult and aged mice (**Figure 1k**). A similar pattern was observed both in whole lysates (**Supplementary Figure 1b-e**) and in PSD-95 immunoprecipitates (**Supplementary Figure 1f-i**).

When analysing human post-mortem tissue at different ages (21 to 89 years old), we found a tendency for a decrease in GluN2B total levels (**Supplementary Figure 1j, k**).

These data demonstrate that NMDARs contribute to synaptic transmission in an age-dependent manner, whereby GluN2B contribution is higher in both infant and aged synapses.

### APP interacts and regulates GluN2B-NMDARs at immature synapses

There are several mechanisms and protein interactors able to regulate NMDAR synaptic transmission and GluN2B/A contribution. The amyloid precursor protein (APP) has emerged as a potential candidate, following reports by our group and others that APP interacts and regulates NMDAR surface levels and currents (Pousinha et al., 2017, Cousins et al., 2009, Hoe et al., 2009).

We detected APP in PSD-enriched fractions at all ages, observing a 5-fold increase in infants compared to adults (n=8), whereas the APP post-synaptic levels remain low in aged synapses (**Figure 2a, b**). In addition, NMDARs co-immunoprecipitated with APP in infant, adult and aged hippocampal synaptosomes. This interaction is mainly established with GluN2B during postnatal development, but it occurs with both GluN2A and GluN2B in adult and aged animals (**Figure 2c**). This interaction was also present in human brain tissue (**Figure 2d**).

**Figure 2 -.**
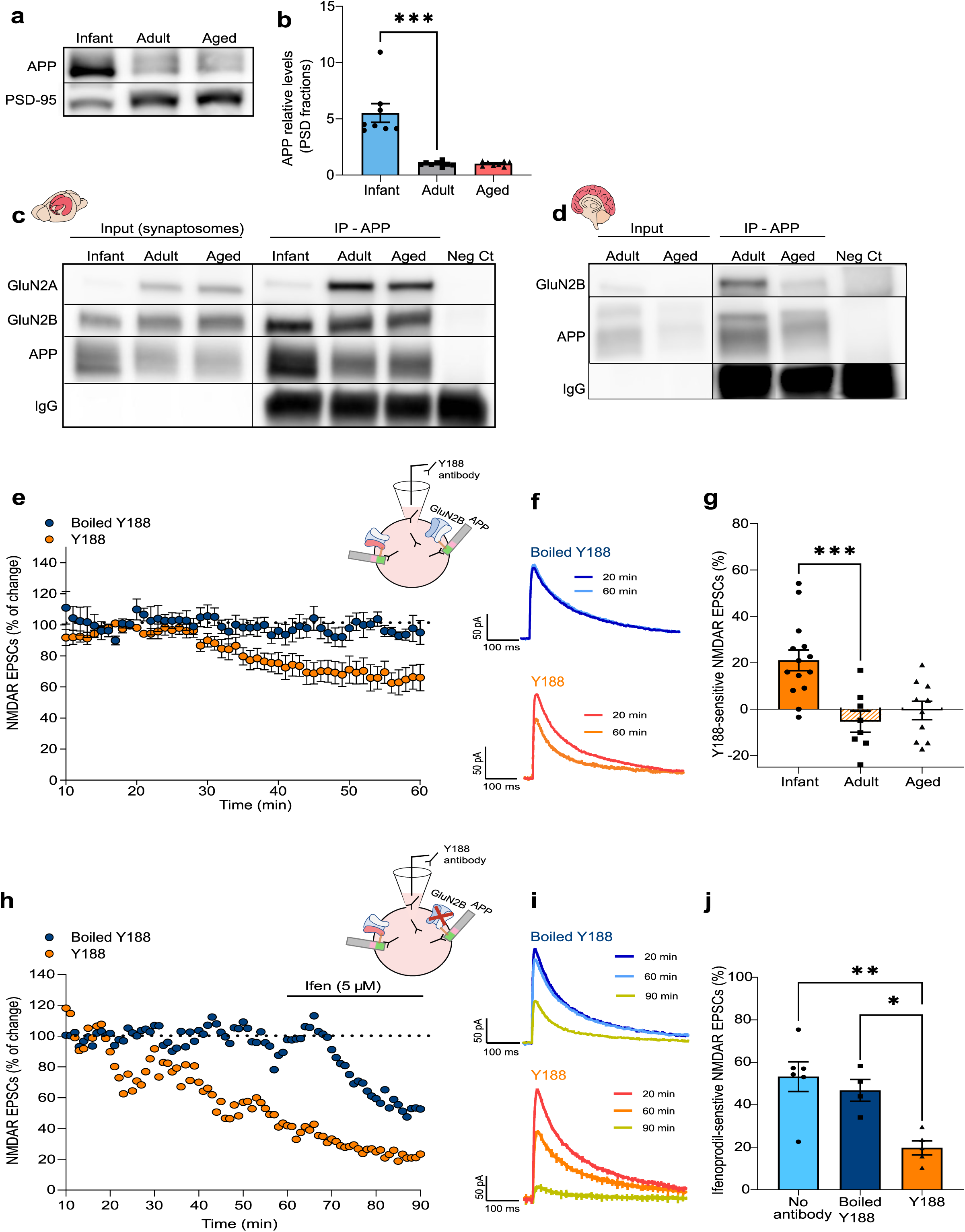
APP interacts and regulates GluN2B-NMDARs at immature synapses. (a) Representative western blot of hippocampal PSD-enriched fractions from infant, adult and aged wild-type C57BL/6 mice. Membranes were immunoblotted with antibodies for APP and PSD-95. (b) APP levels were normalized with PSD-95 and are expressed as the mean ± SEM (Kruskal Wallis test followed by Uncorrected Dunn’s test for multiple comparisons using the adult group as reference, ***p<0.001, n=8). (c) Representative western blot of synaptosome fractions from the hippocampi of infant, adult and aged wild-type C57BL/6 mice immunoprecipitated for APP. Membranes were immunoblotted with antibodies for GluN2A, GluN2B and APP. (d) Representative western blot of postmortem brain tissue (prefrontal cortex) from human subjects (adult=22 and aged=89 years old) immunoprecipitated for APP. Membranes were immunoblotted with antibodies for GluN2B and APP. (e) Time course of NMDAR EPSC amplitude measured by whole-cell patch clamp in CA1 pyramidal neurons of infant C57BL/6 wild-type mice during 60 min of incubation with an antibody against the APP C-terminal (Y188). In the control condition, the antibody was heat inactivated (boiled Y188). Results are expressed as the mean ± SEM (n=6-10). The schematic diagram shows the strategy used to mask the APP C-terminal domain - the antibody was added to the intracellular solution in the patch pipette to diffuse into the intracellular space. (f) Traces show NMDAR EPSCs recorded at 20min (baseline) and 60min. The Y188 antibody was inside the patch pipette during the whole course of the experiment (60min). (g) The percentage of Y188-sensitive NMDAR EPSCs was determined for infant, adult and aged C57BL/6 wild-type mice. The effect was calculated comparing the baseline amplitude (15-20 min) with the final amplitude (60 min) and normalized with the control condition (boiled Y188) for each age. Results are expressed as the mean ± SEM (One-way ANOVA followed by an Uncorrected Fisher’s LSD’s multiple comparisons test, ***p<0.001, n=8-14). (h) Time course of NMDAR EPSC amplitude measured by whole-cell patch clamp in CA1 pyramidal neurons of infant C57BL/6 wild-type mice during 90 min of incubation with the Y188 antibody and perfusion with ifenprodil (5µM) at 60-90 min. In the control condition, the antibody was heat inactivated (boiled Y188) (n=1). The schematic diagram shows the strategy used to block APP (Y188 antibody in the patch pipette) and to inhibit GluN2B-NMDAR (ifenprodil perfusion). (i) Traces show NMDAR EPSCs recorded at 20min (baseline), 60min and 90min. The Y188 antibody was inside the patch pipette during the whole course of the experiment (60min), whereas Ifenprodil perfusion occurred from 60 to 90min. (j) The percentage of ifenprodil-sensitive NMDAR EPSCs in infant mice was calculated comparing the amplitude at 60 min with the final amplitude (90 min). The effect of ifenprodil on NMDAR EPSCs was calculated in neurons without antibody incubation (No antibody, used as reference), incubated with the Y188 antibody (Y188) or the heat-inactivated antibody (boiled Y188) for 90 min. Results are expressed as the mean ± SEM (One-way ANOVA followed by Tukey’s multiple comparisons test, **p<0.01, *p<0.05, n= 4-6).

We previously reported that the loss of GluN2B synaptic contribution following in utero silencing of APP could be reverted by delivering the AICD peptide to the neurons (Pousinha et al., 2017), thus suggesting that APP – NMDR interaction occurs through APP intracellular domain. We therefore interfered with APP – NMDAR interaction while recording NMDAR EPSCs, by introducing an APP C-terminal antibody (clone Y188, 2.5nM) in the recording pipette (**Figure 2e**). We observed a reduction of 21,11 ± 4,47% (mean ± SEM) in NMDAR EPSCs in infant mice (n=14), compared to the control condition (boiled Y188 antibody) (as illustrated in **Figures 2e-g**). This effect was not stronger by increasing the Y188 antibody concentration to 5nM (**Supplementary Figure 2**). To test if this response is mediated by GluN2B-NMDARs, we perfused hippocampal slices with a GluN2B selective antagonist, ifenprodil (5µM), for 30 min at the end of the experiment (60-90 min) (**Figure 2h**). As shown, the effect of ifenprodil was reduced in neurons previously incubated with the Y188 antibody, when comparing to the control condition (19,76 ± 3,292 vs. 46,82 ± 5,160%, mean ± SEM, n=4-6) (**Figure 2h-j**). This data indicates that interfering with the APP intracellular domain in infant mice relates to a decrease in GluN2B synaptic contribution. In contrast, we found no alterations in NMDAR-mediated currents in adult and aged mice by blocking APP (n=8-10) (**Figure 2g**).

In conclusion, these results show that APP interacts with GluN2B at immature synapses, thus regulating their contribution to NMDAR EPSCs.

### APP modulates the GluN2B-NMDAR synaptic content in immature synapses

The reported impact of APP on NMDAR transmission suggested a role for APP in regulating NMDARs during postnatal development. To assess the outcomes of APP depletion in immature neurons, we transfected primary neuronal cultures (7 days *in vitro*) with a plasmid encoding a short-hairpin RNA (shRNA) sequence against APP (shAPP) or the respective control (shRNA with no silencing effect, shCTR). The APP knockdown was efficient, leading to a reduction of approximately 80% in APP immunoreactivity, when comparing to the control condition (n=20-21 cells, 3 independent cultures), evaluated 7 days post-transfection (14 days *in vitro*) (**Figure 3a, b**).

**Figure 3 -.**
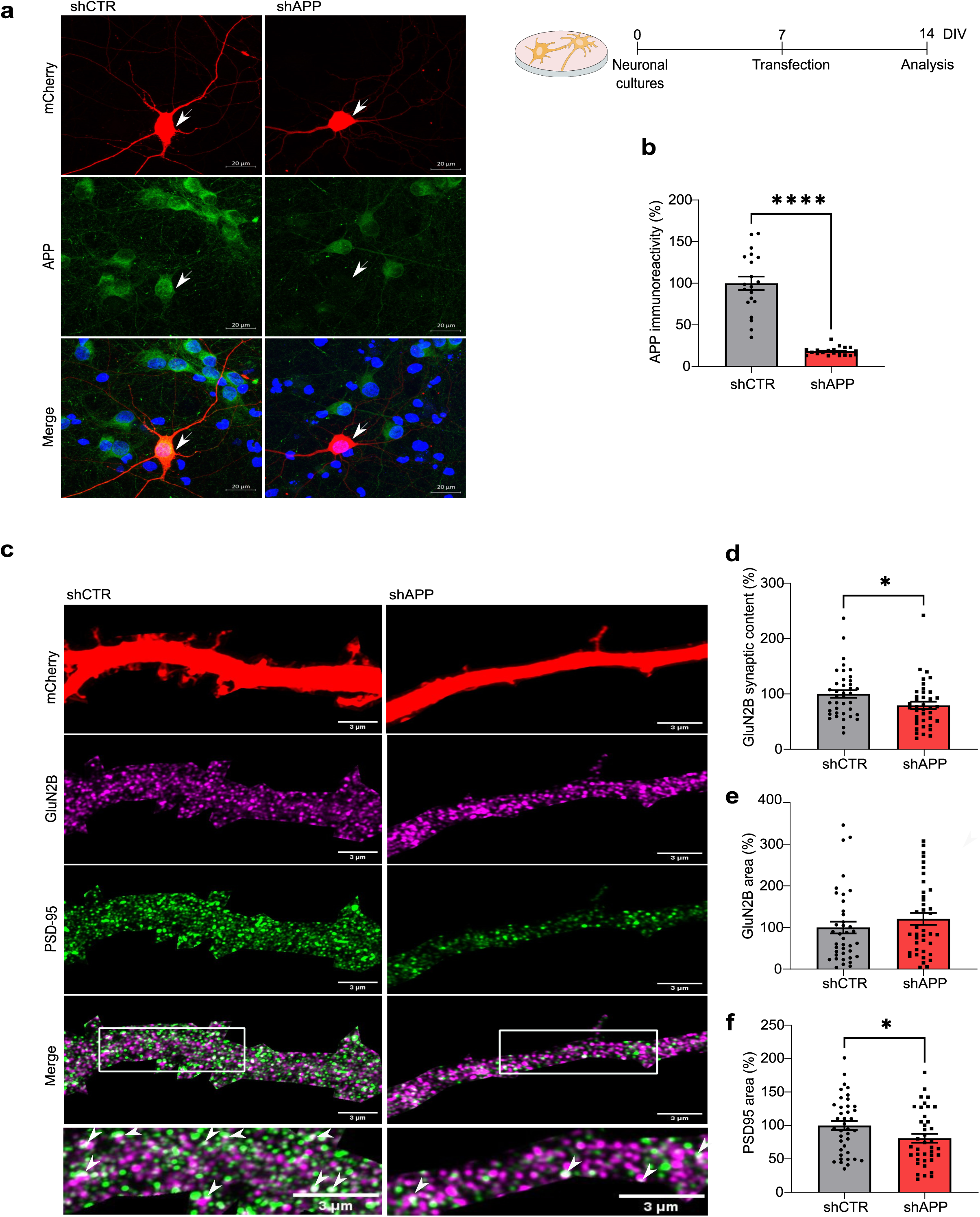
APP modulates the GluN2B-NMDAR synaptic content in immature neurons. (a) Representative immunocytochemistry analysis of APP immunofluorescence in rat primary neuronal cultures (14 days *in vitro* (DIV)) transfected with shAPP or the respective control (shCTR) at DIV7, as indicated in the timeline. mCherry (reporter plasmid) is shown in red, APP is labelled in green and cell nuclei are stained with Hoechst in blue. Transfected neurons are indicated by arrows. (b) APP immunoreactivity (%) in transfected neurons is expressed as the mean ± SEM, using the control condition as reference (Mann-Whitney test, ****p<0.0001, n=20-21 cells, 3 independent cultures). (c) Representative immunocytochemistry analysis of rat primary neuronal cultures (DIV14) transfected with shAPP or the respective control (shCTR) at DIV7. mCherry (reporter plasmid), labelled in red, was used to identify dendrites of transfected neurons. GluN2B is shown in magenta and PSD-95 is labelled in green. Higher magnification images are shown at the bottom, with arrows indicating GluN2B/PSD95 co-localization. (d, e) Results of analysis of GluN2B synaptic content (GluN2B – PSD 95 co-localization) and area from dendrites of transfected neurons are expressed as the mean ± SEM, using the control condition as reference (%) (Mann-Whitney test, *p<0.05, n= 39, 3 independent cultures) (f) Results of analysis of PSD95 area from dendrites of transfected neurons are expressed as the mean ± SEM, using the control condition as reference (%) (Unpaired t-test, *p<0.05, n= 39, 3 independent cultures).

APP depleted neurons showed a significant reduction in the percentage of GluN2B clusters that co-localize with PSD-95 (post-synaptic marker) (approximately 20%) (**Figure 3c, d**). The GluN2B relative dendritic area, average particle size and fluorescent density were not altered by APP depletion, indicating that GluN2B total levels/area and clustering remained constant (n=39 dendrites, 3 independent cultures) (**Figure 3c, e; Supplementary Figure 3a, b**).

Considering the impact of GluN2B on synaptic maturation, in which PSD-95 recruitment is a crucial step (Elias et al., 2008), we also evaluated if this process is impaired in APP-depleted neurons. We found a reduction in PSD-95 dendritic area and average particle size (**Figure 3c, f; Supplementary Figure 3c**), whereas the fluorescence density was not altered (n=39 dendrites, 3 independent cultures) (**Supplementary Figure 3d**).

These findings show that APP regulates GluN2B synaptic content and PSD-95 clustering in immature neurons.

### Age-related increase in APP processing contributes to higher GluN2B synaptic contribution

Since the increase in GluN2B contribution to NMDAR EPSCs in aged mice is not correlated with APP post-synaptic levels or APP-NMDAR interaction, we hypothesized that APP in its full-length form is not responsible for these alterations. In fact, we had previously reported that an APP-derived fragment, the AICD, has the ability to regulate synaptic GluN2B in CA1 pyramidal neurons (Pousinha et al., 2017).

We thus characterized APP processing throughout time in the hippocampus of wild-type C57BL/6 mice by detecting APP-full length (APP), APP C-terminal fragments (APP CTFs derived from α- (non-amyloidogenic) and β-secretase (amyloidogenic) pathways) and AICD (**Figure 4a**). When compared to adults, aged mice displayed no alterations in the APP levels (**Figures 4a, b**), but instead they exhibited an increase in APP processing. Accordingly, the ratio between APP CTFs and APP exhibits an approximate 1,5-fold increase in aged mice, when compared to the reference values for adults (**Figure 4a, c**). Of note, we could detect a tendency to increase of AICD fragment levels (**Figure 4a, d**), even if it is well known that AICD is difficult to detect biochemically and unstable (Pimplikar & Suryanarayana, 2011). We used a triple transgenic mouse (3xTg-AD) as a positive control, since this model exhibits a marked increase in APP-derived fragments with a predominant accumulation of CTF-β over the smaller CTF-α (Lauritzen et al., 2012) (**Figure 4a**).

**Figure 4 -.**
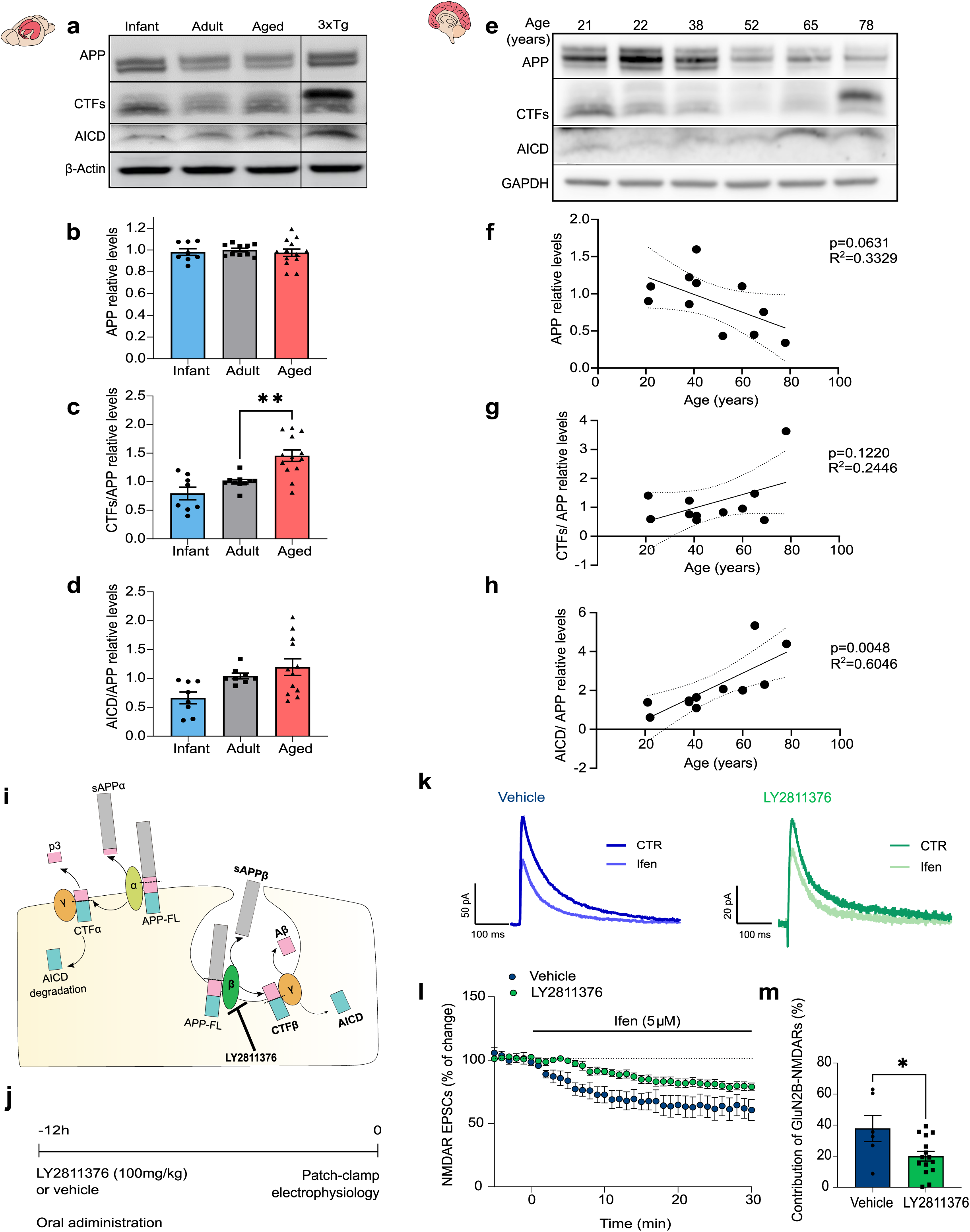
Age-related increase in APP processing contributes to higher GluN2B synaptic contribution. (a) Representative western blot of hippocampal lysates from infant, adult and aged C57BL/6 wild-type mice. Membranes were immunoblotted with antibodies for APP C-terminal (to detect APP (APP), APP C-terminal fragments (CTFs) and the APP Intracellular Domain (AICD)) and β-actin. A female triple transgenic mouse (3xTg, 6 months) was used as a positive control for APP-derived fragments. (b) Results from blots shown in (a) from mouse hippocampal lysates were normalized with β-actin and are expressed as the mean ± SEM (One-way ANOVA followed by an Uncorrected Fisher’s LSD’s multiple comparisons test using the adult group as reference, n=8-13) (c) Results from blots shown in (a) from mouse hippocampal lysates show the ratio between CTFs and APP (Kruskal Wallis test followed by Uncorrected Dunn’s test using the adult group as reference, **p<0.01, n=8-13). (d) Results from blots shown in (a) from mouse hippocampal lysates show the ratio between AICD and APP (One-way ANOVA followed by an Uncorrected Fisher’s LSD’s multiple comparisons test using the adult group as reference, n=8-13). (e) Representative western blot of prefrontal cortex human samples (21 to 78 years old). Membranes were immunoblotted with antibodies for APP C-terminal (to detect APP, CTFs and AICD) and GAPDH. (f) Linear regression graph calculated from blots as shown in (e) shows the variation in APP relative levels (normalized to GAPDH) depending on the age of human subjects (n=11). Statistical analysis was performed using Pearson’s correlation (two-tailed p value). Dotted lines represent the 95% confidence intervals. The values obtained for 20–25-year-old subjects were used as reference. (g) Linear regression graph calculated from blots as shown in (e) shows the variation in the ratio between APP CTFs and APP depending on the age of human subjects (n=11). Statistical analysis was performed using Pearson’s correlation (two-tailed p value). Dotted lines represent the 95% confidence intervals. The values obtained for 20–25-year-old subjects were used as reference. (h) Linear regression graph calculated from blots as shown in (e) shows the variation in the ratio between AICD and APP depending on the age of human subjects (n=11). Statistical analysis was performed using Pearson’s correlation (two-tailed p value). Dotted lines represent the 95% confidence intervals. The values obtained for 20–25-year-old subjects were used as reference. (i) Schematic diagram shows the LY2811376 effect on APP processing. Inhibition of the β-secretase, it interferes with the APP amyloidogenic processing pathway, but not with the non-amyloidogenic pathway (α-secretase). (j) Representation of the timeline for the β-secretase inhibitor (LY2811376) treatment in aged mice. (k) Traces show NMDAR EPSCs recorded before (CTR) and after 30 min of Ifenprodil 5 µM perfusion (Ifen). (l) Time course of ifenprodil (5µM) effect on pharmacologically isolated NMDAR EPSC amplitude in CA1 pyramidal neurons, measured by whole-cell patch clamp in aged C57BL/6 wild-type mice (vehicle vs. LY2811376, administration 12h prior to patch-clamp recordings). Results are expressed as the mean ± SEM (n=6-15). (m) GluN2B contribution was calculated as the percentage of change in NMDAR EPSCs after ifenprodil perfusion (for 30 min) in aged C57BL/6 wild-type mice (vehicle vs. LY2811376). Results are expressed as the mean ± SEM (Unpaired t-test, *p<0.05).

To elucidate whether this age-related pattern of APP processing is also observed in the human brain, we prepared lysates from human post-mortem brain tissue (prefrontal cortex) of subjects with different ages (21 to 78 years old) (**Figure 4e**). We found a negative correlation between age and APP levels (**Figure 4e, f**). Instead, a positive correlation seems to occur regarding APP processing into CTFs and AICD with age (**Figure 4e, g, h**). We detected a similar pattern of APP levels/processing in human AD patients (n=4, 65-81 years old) compared to aged controls (n=4, 65-89 years old) (**Supplementary Figure 4**), albeit more variable.

The aging impact on APP processing is likely independent of cellular APP levels alterations, which we found unaltered. APP endocytosis was shown to be upregulated in aging, thus favouring APP processing through the amyloidogenic pathway (Burrinha et al., 2021). We therefore blocked APP amyloidogenic processing in aged mice by oral administration of a β-secretase 1 (BACE 1) (BI) inhibitor (LY2811376; 100mg/kg), 12 h prior to patch-clamp recordings (**Figure 4i, j**). The aged mice treated with LY2811376 showed a significant decrease in GluN2B synaptic contribution when compared to the vehicle-treated group (20,07% ± 3,118 vs. 37,95% ± 8,445, mean ± SEM) (**Figure 4k-m**), which is similar to levels found in untreated adult mice.

We concluded that the observed increased APP processing upon aging, detected both in human and mouse brain, contributes to the accumulation of APP-derived fragments, leading to a significant increase in GluN2B synaptic contribution.

## Discussion

We show that synaptic GluN2B-NMDARs are regulated by APP in an age-dependent manner. During postnatal development, APP interacts with GluN2B at the synapse and modulates its synaptic content and evoked currents, possibly having an impact on synaptic maturation. On the other hand, APP-derived amyloidogenic fragments contribute to increased GluN2B synaptic contribution upon aging, potentially underlying age-related synaptic impairments.

We found that the APP interaction regulates the GluN2B-NMDARs function during development, a life stage when the GluN2B subtype predominates, accounting for more than 50% of NMDAR EPSCs in the CA1 hippocampal region. The electrophysiological output reflected the synaptic NMDAR subunit composition, since infant mice exhibited a high GluN2B/GluN2A ratio in protein levels, according to the previously described NMDAR developmental switch (GluN2B to GluN2A) that occurs during postnatal development (Williams et al., 1993). These data indicate a crucial role for APP at the post-synapse, in contrast with most of the studies so far, which focused on APP function at the pre-synapse (Marquez-Sterling et al., 1997, Lyckman et al., 1998), given the low APP levels detected in PSD-enriched fractions from the adult brain (Kim et al., 1995).

We now found that APP subcellular localization is age dependent, since we could observe abundant APP at the post-synapse in infant mice, whereas the levels decline in adulthood and aging, thus suggesting that the role of APP at the synapse shifts with age. Indeed, we proved that the presence of a monoclonal antibody binding to the APP C-terminal interferes with NMDAR-evoked currents during postnatal development. Importantly, this experimental strategy allows to elucidate the role of APP at the post-synapse without interfering with the pre-synaptic compartment. The NMDAR-APP interaction was further investigated in rat hippocampal primary neuronal cultures at immature state (DIV 7-14), since this model recapitulates the synapse development process (Grabrucker et al., 2009) and the NMDAR developmental switch (Corbel et al., 2015), while providing appropriate imaging resolution to study synaptic proteins. APP-depleted neurons exhibited a decreased GluN2B synaptic content, thereby confirming that APP is crucial for NMDAR synaptic levels/activity in immature synapses. The mechanism herein described justifies the loss of GluN2B synaptic contribution that we previously observed in infant mice following in utero silencing of APP (Pousinha et al., 2017). Considering the key role of GluN2B currents for synaptic maturation (Durand et al., 1996, Gray et al., 2011, Adesnik et al., 2008), we hypothesize that this process is essential to achieve functional mature synapses during development. This is consistent with our observations in APP-depleted neurons, in which PSD-95 dendritic area and cluster size are reduced. These data might seem contradictory with the observation that the APP KO model only presents synaptic deficits after 10 months of age (Dawson et al., 1999, Tyan et al., 2012). This may result from the functional redundancy of APP and APP-like proteins (APLP1 and ALPL2). In accordance, constitutive triple knockout mice (TKO) die after birth (Heber et al., 2000), whereas Nex-Cre cTKO (conditional triple KO in excitatory forebrain neurons starting during embryonic development) present gross brain morphology alterations (Steubler et al., 2021), showing a crucial role for APP family members during development. Our combination of *in vitro* silencing and acute interference of the APP C-terminal domain *ex vivo*, overcome a possible compensation by APP family members, while allowing to study APP specifically during postnatal developmental stages.

Considering that the antibody against the APP C-terminal had a significant effect on NMDAR-mediated currents in infant mice, this seems to be the domain responsible for interacting and regulating NMDARs at this stage. In particular, the YENPTY motif is highly conserved in APP family members and different species (Shariati & de Strooper, 2013) and known to interact with several proteins (van der Kant & Goldstein, 2015), possibly acting as the interacting site for NMDARs either directly or indirectly. We had previously shown that neurons expressing a mutated form of AICD in the YENPTY motif lose their ability to modulate NMDA currents in the adult rat (Pousinha et al., 2017). Accordingly, mouse models in which the APP C-terminal domain is mutated (Matrone et al., 2012) or depleted (APPdCT15 knock-in mice/ APLP2 KO) (Klevanski et al., 2015) show high postnatal lethality, impairments in synaptic plasticity and memory in adult stages, which might be explained by NMDAR dysregulation, although this hypothesis has not been explored so far.

We could not affect NMDAR EPSCs in adult/aged animals when interfering with the APP C-terminal domain in a 60 min time window, which may allow for different interpretations. Considering the technical approach we used, it is possible that the accessibility of the APP C-terminal epitope at the post-synapse might be precluded in adult stages, considering the extensive alterations that occur in PSD structure and composition after postnatal development (Gonzalez-Lozano et al., 2016). However, we can also postulate that the APP regulation of NMDARs has indeed a higher impact during development, which is consistent with the marked decline of postsynaptic APP levels observed in adult stages. Moreover, it is possible that the APP-NMDAR regulation mainly occurs through GluN2B, whose synaptic contribution declines after development. Our observation of a reduced effect of ifenprodil in APP C-terminal antibody exposed neurons from infant mice shows that GluN2B-NMDARs are highly affected by disruption of APP C-terminal function. Since APP interacts with both GluN2B and GluN2A in adult stages, it is extremely difficult to determine if there is a subunit preferential binding, especially considering that tri-heteromeric complexes (GluN1/GluN2A/GluN2B) also exist in the hippocampus (Rauner & Köhr, 2011). Given the increased association of PSD-95 with NMDARs in mature synapses (Petralia et al., 2005, Elias et al., 2008), we can postulate that the NMDAR synaptic clustering becomes APP-independent upon adulthood. However, the fact that the Camk2a-Cre cTKO mouse model (triple conditional knock-out for APP family members in excitatory forebrain neurons starting at postnatal weeks 3-4) exhibits LTP impairments and reduced NMDAR-mediated responses in adult stages, suggests that some form of NMDAR regulation by APP family members still occurs later in life (S. H. Lee et al., 2020).

We observed significant alterations in NMDAR levels and properties in aged mice, including a decline in GluN2B total levels which is consistent with other studies (Magnusson, 2000, Zhao et al., 2009, Clayton & Browning, 2001, Sonntag et al., 2000, Bai et al., 2004) and potentially contributes to a reduction in NMDAR agonist binding, as previously reported (Magnusson, 2000, Piggott et al., 1992). We also found that NMDAR deactivation kinetics become slower upon aging, possibly explaining the increased duration of NMDAR field excitatory postsynaptic potentials (fEPSPs) previously described in aged rodents (Jouvenceau et al., 1998). This data correlates with the augmented GluN2B contribution we observed in aged mice, possibly reflecting the higher GluN2B synaptic retention reported in previous studies (Zamzow et al., 2013, al Abed et al., 2020). We postulate that APP contributes to these alterations, controlling NMDAR function in aged synapses. This hypothesis is supported by the synaptic plasticity and learning/memory deficits observed in APP KO aged mice, but not in earlier stages (10 months vs. 4 months) (Dawson et al., 1999, Tyan et al., 2012).

Our findings suggest that the APP-NMDAR regulation in aged synapses occurs through APP-derived fragments rather than through the full-length protein. We detected a significant enhancement of APP processing in aged mice, leading to an accumulation of APP CTFs. Importantly, we observed the same profile in human brain samples, in which we established a positive correlation between APP processing and aging. Considering the previously described accumulation of CTFs (Burrinha et al., 2021) and increased BACE1 activity (Fukumoto et al., 2004) in aged mice, we postulate that APP-derived amyloidogenic fragments accumulate and alter NMDAR function. To test this hypothesis, we used a β-secretase 1 (BACE 1) inhibitor in aged animals to block the APP amyloidogenic processing, which resulted in a decrease in GluN2B synaptic contribution of about 20%, getting closer to the results obtained in adult mice. The selected dosage and timing of administration of BACE 1 inhibitor does not affect the non-amyloidogenic processing pathway (Filser et al., 2015) (May et al., 2011), thus excluding putative protective roles of soluble APP alpha fragments (Müller et al., 2017).

Although we did not single out the APP amyloidogenic fragment which is responsible for this effect, the AICD has emerged as a potential target, since we have previously shown that this peptide increases GluN2B synaptic contribution in adult synapses (Pousinha et al., 2017) and though not statistically significant, we could observe a tendency to AICD/APP increased levels. Moreover, BACE1 inhibition is known to interfere with AICD signalling, since the biologically active AICD fragment mainly derives from the amyloidogenic pathway (Edbauer et al., 2002, Goodger et al., 2009). Considering that an incubation with the APP C-terminal antibody (60 min) did not alter the effects we have seen in aging, we postulate that a more prolonged mechanism, possibly involving transcriptional targets, is occurring at this age. Since the ratio of GluN2B/GluN2A protein levels was not directly affected, possibly alternative transcriptional targets are involved, such as kinases/phosphatases that are known to regulate GluN2B phosphorylation status, Ca^2+^ permeability (Murphy et al., 2014) or trafficking. It is possible that these functional alterations arise at a first stage as a compensatory mechanism against the decline in GluN2B levels that occurs upon aging. However, the maintenance of this response may become toxic to neurons, considering that GluN2B-NMDAR carry more Ca^2+^ per unit of current, leading to calcium dyshomeostasis (Kumar et al., 2009).

Therefore, our findings could help to clarify an apparent paradox in the field: although NMDAR levels are known to decline in physiological aging and Alzheimer’s Disease (Hynd et al., 2004b, Hynd et al., 2004a, Jacob et al., 2007, Mishizen-Eberz et al., 2004), NMDAR antagonists such as memantine show beneficial effects in both conditions (Norris & Foster, 1999, Pietá Dias et al., 2007, Beracochea et al., 2008, Reisberg et al., 2003). Overall, we show that the NMDARs that remain at aged synapses have different properties than in adulthood, staying open for longer times and being more dependent on the GluN2B subtype. An imbalance in subunit contribution towards GluN2B is expected to decrease the threshold for LTP (Yashiro & Philpot, 2008), possibly contributing to the LTD-LTP shift reported by our group in aged rats (Temido-Ferreira et al., 2020). Therefore, the increased contribution of GluN2B may lead to synaptic dysregulation in physiological aging, while increasing the susceptibility to neurodegeneration, in which GluN2B-NMDARs become overactivated (Li et al., 2011). This mechanism might be particularly relevant in AD, since the pathological accumulation of APP amyloidogenic fragments is expected to further enhance GluN2B contribution. Accordingly, memantine is an approved drug for moderate to serious forms of AD (Reisberg et al., 2003).

In conclusion, we describe two alternative mechanisms by which APP controls GluN2B-NMDARs, depending on the age. We have discovered a new physiological role for APP at the post-synapse, being essential to maintain GluN2B synaptic content/currents in immature synapses. Moreover, we show that the age-related increase in APP processing contributes to a higher GluN2B synaptic contribution. While the first mechanism might play a crucial role in synaptic maturation, the latter could be involved in age-associated synaptic dysfunction.

## Methods

### Human samples

The use of human samples was conducted in accordance with the Helsinki Declaration as well as national ethical guidelines. Protocols were approved by the Local Ethics Committee and the National Data Protection Committee. Samples of post-mortem brain tissue from the prefrontal cortex were obtained from the National Institute of Legal Medicine and Forensic Sciences, Coimbra, Portugal or Neuro-CEB Biological Resource Center (BRC), France. Samples from human AD patients correspond to Braak Stage VI.

### Animals

Animal procedures were performed at the Rodent Facility of Instituto de Medicina Molecular, that is a licensed establishment (license number 017918/2021) in compliance with the European Directive 2010/63/EU, transposed to Portuguese legislation in DL 133/2013. All animal research projects carried out at iMM are reviewed by the Animal Welfare Body (ORBEA-iMM) to ensure that the use of animals is carried out in accordance with the applicable legislation and following the 3R’s principle. Environmental conditions were kept constant: food and water ad libitum, 22-24 °C, 45-65% relative humidity, 14h light/10 dark cycles, 3 to 4 mice per cage.

Experiments performed at IPMC were done according to policies on the care and use of laboratory animals of European Communities Council Directive (2010/63) and the protocols were approved by the French Research Ministry following evaluation by a specialized ethics committee (protocol number 00973.02) All efforts were made to minimize animal suffering and reduce the number of animals used. The animals were housed three per cage under controlled laboratory conditions with food and water ad libitum, a 12 hr dark light cycle and a temperature of 22 ±2°C.

Male and female wild-type C57BL/6 mice at different ages were used: infant (7-10 days), adult (10 – 16 weeks) and aged (18 – 20 months). A C7BL/6-129SvJ female mouse bearing three mutations (3xTg-AD) associated with familial AD (amyloid precursor protein [APPswe], presenilin-1 [PSEN1] and microtubule-associated protein tau [MAPT]) (Mutant Mouse Research and Resource Center at The Jackson Laboratory) was used as control.

### BACE1 Inhibitor

LY2811376 was obtained from Medchem Express (Sweden) and prepared in 10% DMSO, 40% PEG300, 5% Tween-80, 45% saline. C57BL/6 aged mice (18 – 20 months) received LY2811376 at 100 mg/kg body weight by oral gavage as described in (Filser et al., 2015). Animals treated with LY2811376 (n=3) or vehicle (n=2) were sacrificed ≈12h after treatment.

### Patch Clamp electrophysiology

Newborn mice were anesthetised through hypothermia, whereas adult and aged mice were anesthetized [ketamine (150 mg/kg)/xylazine (10 mg/kg)]. All groups were transcardially perfused with artificial cerebrospinal fluid (aCSF) for slice preparation. Acute transverse hippocampal slices (250 µm) from wild-type C57BL/6 mice were prepared on a vibratome (Microm HM600V, Thermo Scientific, France) in ice-cold dissecting solution containing (in mM): 234 sucrose, 2.5 KCl, 0.5 CaCl2, 10 MgCl_2_, 26 NaHCO3, 1.25 NaH_2_PO_4_ and 11 D-glucose, oxygenated with 95% O_2_ and 5% CO_2_, pH 7.4. Slices were incubated for 60 min at 37°C, in an artificial CSF (aCSF) solution containing (in mM): 119 NaCl, 2.5 KCl, 1.25 NaH_2_PO_4_, 26 NaHCO_3_, 1.3 MgSO_4_, 2.5 CaCl_2_ and 11 D-glucose, oxygenated with 95% O2 and 5% CO2, pH 7.4. Slices were used after recovering for another 30 min at room temperature. To measure pharmacologically isolated NMDAR EPSCs, slices were perfused with the oxygenated aCSF at 31 ± 1°C in the continuous presence of 50 mM picrotoxin (Sigma-Aldrich, dissolved in DMSO) to block GABAergic transmission and DNQX (10 µM) to block AMPA receptors.

Recording pipettes (5-6 MΩ) for voltage-clamp experiments were filled with a solution containing the following: 117.5 mM Cs-gluconate, 15.5 mM CsCl, 10 mM TEACl, 8 mM NaCl, 10 HEPES, 0.25 mM EGTA, 4 mM MgATP and 0.3 NaGTP (pH 7.3; osmolarity 290-300 mOsm). Slices were visualized on an upright microscope with IR-DIC illumination (Scientifica, Ltd). Whole-cell recordings were performed using a Multiclamp 700B (Molecular Devices) amplifier, under the control of pClamp10 software (RRID:SCR_011323) (Molecular Devices). The Schaffer collateral pathway was stimulated at 0.10 Hz using electrodes (glass pipettes filled with aCSF) placed in the stratum radiatum.

After a tight seal (>1 GW) on the cell body of the selected neuron was obtained, whole-cell patch clamp configuration was established, and cells were left to stabilize for 2–3 min before recordings began. Pharmacologically isolated NMDAR EPSCs were recorded from cells voltage clamped at +40 mV. Holding current and series resistance were continuously monitored throughout the experiment, and if either of these two parameters varied by more than 20%, the experiment was discarded. Electrical stimulation was adjusted to elicit EPSCs of approximately 150 pA amplitude in the different studied groups.

For the analysis of NMDAR EPSCs deactivation kinetics, decay time was fitted with a double exponential function, using Clampfit software, to calculate both slow and fast decay time constants, τ_fast_ and τ_slow_, respectively. The weighted time constant (τ_weighted_) was calculated using the relative contribution from each of these components, applying the formula: τ_w_ = [(a_f_. τ_f_) + (a_s_. τ_s_)]/(a_f_ + a_s_), where a_f_ and a_s_ are the relative amplitudes of the two exponential components, and τ_f_ and τ_s_ are the corresponding time constants.

To calculate GluN2B-NMDAR contribution, NMDAR EPSCs were measured immediately before (5 min) and 25-20 min after ifenprodil (5µM) perfusion to selectively block GluN2B-NMDARs.

To interfere with the APP – NMDAR interaction, the APP C-terminal antibody Y188 (ab32136, Abcam) was added to the intracellular solution to a final concentration of 0.4µg/mL (2.57nM). In the control condition, the same antibody was heat-inactivated by incubation at 98°C for 10min.The percentage of reduction in NMDAR EPSCs due to the APP C-terminal antibody incubation was calculated comparing the baseline amplitude (15-20min) with the final amplitude (55-60 min) and normalized with the control condition for each age.

### Protein analysis

#### Fractionation into PSD-enriched fractions

Hippocampi from C57BL/6 wild-type mice were dissected and snap-frozen in liquid nitrogen. The fractionation protocol was adapted from (Frandemiche et al., 2014), all centrifugation steps were performed at 4°C and all solutions contained protease/phosphatase inhibitors. Samples were first homogenized using a Glass/Teflon Potter Elvehjem homogenizer in Buffer I (0.32M sucrose and 10mM HEPES, pH 7.4) and then centrifugated (1.000 g for 10min) to remove nuclei and cell debris. This centrifugation step was repeated 3 times, until the supernatant was completely clear and finally subjected to a centrifugation at 12.000 g for 20min. The pellet was recovered, resuspended in Buffer II (4mM HEPES and 1 mM EDTA, pH 7.4) and centrifuged twice at 12.000 g for 20 min. The pellet was then resuspended in 25µL of Buffer III (20 mM HEPES, 100mM NaCl, 0,5% Triton X100, pH= 7.2) and incubated 1h at 4°C with mild agitation. By centrifuging the samples at 12.000 g for 20 min it was possible to pellet the synaptosome membrane fraction, whereas the supernatant was collected as the non-postsynaptic density membrane fraction (non-PSD) or Triton-soluble fraction. Finally, the pellet was solubilized in 25µL Buffer IV (20 mM HEPES, 0.15 mM NaCl, 1% TritonX-100, 1% sodium deoxycholate, 1% SDS, pH 7.5) for 1 h at 4°C and centrifuged at 10.000 g for 15 min. The supernatant contained the PSD or Triton-insoluble fraction. The integrity of non-PSD was verified by immunoblotting for synaptophysin, which was enriched in the non-PSD fraction, and the integrity of the PSD fraction was confirmed by the immunoblotting of PSD-95 enriched in this compartment.

### Synaptosomes preparation

The protocol for synaptosome preparation was adapted from (Lopes et al., 1999), all centrifugation steps were performed at 4°C and all solutions contained protease/phosphatase inhibitors. Hippocampi from C57BL/6 wild-type mice were dissected, resuspended in a 0.32 M sucrose solution containing 50mM Tris, 2 mM EDTA, pH=7.6 and homogenized using a Glass/Teflon Potter Elvehjem homogenizer. The suspension was centrifuged at 3.000 g during 10 min. The supernatant was collected and centrifuged at 1.000 g for 12 min. The pellet was resuspended in 1.8 ml of a 45% vol/vol Percoll solution made up in a Krebs-Ringer solution (140mM NaCl, 1mM EDTA, 10mM HEPES, 5mM KCl, pH= 7.4). After centrifugation at 21.100g for 2 min, the top layer was collected (synaptosome fraction) and washed twice in Krebs-Ringer solution (centrifugation at 21.100g, 2 min). When synaptosome fractions were used for co-immunoprecipitation experiments, they were subsequently resuspended in the respective buffer (50mM Tris HCl pH 7.5; 150 mM NaCl; 2 mM EDTA; 1% Triton with protease and phosphatase inhibitors).

### Co-immunoprecipitation

For co-immunoprecipitation from total lysates, hippocampi from C57BL/6 wild-type mice were dissected and snap-frozen in liquid nitrogen. Frozen tissue was resuspended in immunoprecipitation buffer (50mM Tris HCl pH 7.5; 150mM NaCl; 2mM EDTA; 1% Triton with protease and phosphatase inhibitors) and homogenized using a Glass/Teflon Potter Elvehjem homogenizer. Following a centrifugation at 1.000g, 10 min 4°C the supernatant was collected. Protein quantification of total lysates and synaptosome fractions was performed using the BioRad DC Protein assay kit. The immunoprecipitation protocol was adapted from (Tomé et al., 2021). For each sample, 50µL of Dynabeads were washed 3 times with washing buffer (0.1% BSA; 2mM EDTA in PBS). Dynabeads were then resuspended in 500uL washing buffer and the appropriate volume of antibody: APP C-terminal Y188 (3µg, ab32136, Abcam), PSD95 (3µg, ab18258, Abcam) or Normal rabbit IgG (3µg, 12-370, Merck Millipore) and incubated overnight at 4°C under rotation. Following 3 washing steps with washing buffer, dynabeads were resuspended in 350uL washing buffer and incubated with 500µg of protein lysate diluted in Immunoprecipitation buffer (500µL) for 2h at 4°C under rotation. Following 5 washing steps with Immunoprecipitation buffer, dynabeads were gently resuspended in 60µL of pre-heated 2x sample buffer (140mM Tris HCl pH 6.8, 4% SDS, 13.6% glycerol, 272mM DTT, 0.004% Blue bromophenol) in RIPA (50mM Tris, 1mM EDTA, 150mM NaCl, 0.1% SDS, 1%Tergitol-type NP-40, pH 8.0). Finally, samples were incubated for 10 min at 95°C, the supernatant was collected and used for Western Blot analysis.

### Western blotting

For the preparation of total protein lysates, hippocampi from C57BL/6 wild-type mice were dissected and snap-frozen in liquid nitrogen. Mouse and human frozen tissue samples were resuspended in A-EDTA buffer (10 mM HEPES-KOH pH 7.9, 10 mM KCl, 1.5 mM MgCl_2_, 0.1 mM EDTA, 0.3% NP-40 with protease and phosphatase inhibitors) and homogenized using a Glass/Teflon Potter Elvehjem homogenizer, as described in (Pousinha et al., 2017). Following protein quantification using BioRad DC Protein assay kit, lysates were diluted in water and 5x sample buffer (Final concentration: 50mM Tris HCl pH 6.8, 2% SDS, 6% glycerol, 0.1% bromophenol blue, 121mM DTT) and denatured at 95°C for 10 min.

For APP, APP-CTFs and AICD analysis, optimal conditions for low molecular mass proteins separation were used, as described in (Willem et al., 2015). Proteins were separated using precast gradient Tricine Protein Gels (10–20%, 1 mm, Novex) in Tris-tricine buffer (1M Tris, 1M Tricine, 1% SDS). Samples were electro-transferred at 400mA for 1h to 0.2µm Nitrocellulose membranes using a Tris Glycine buffer (25 mM Tris, 190 mM glycine) with 20% ethanol. Proteins transferred to nitrocellulose membranes were additionally denatured by boiling the membrane in PBS for 5 min, acting as an antigen retrieval step to detect AICD, as described in (Pimplikar & Suryanarayana, 2011).

For all the remaining proteins, electrophoresis was performed in Tris-glycine buffer with 10% SDS using 10-12% and 4% acrylamide resolving and stacking gels, respectively. Proteins were electro-transferred to 0.45µm Polyvinylidene fluoride (PVDF) membranes in Tris-glycine buffer with 20% methanol at 350mA for 90 min.

After transfer, all membranes were blocked with 3% BSA in TBS-T (20 mM Tris,150 mM NaCl, 0.1% Tween-20) at room temperature (RT) for 1h and incubated with primary antibodies (diluted in 3% BSA TBS-T) overnight at 4°C. The following antibodies were used: APP C-terminal Y188 (1:1000, ab32136, Abcam), GluN2A (1:200, sc-136004, Santa Cruz), GluN2B (1:1000, D15B3, Cell Signalling), PSD95 (1:1000, D27E11, Cell signalling), Synaptophysin (1:200, S5768, Merck Millipore), β-actin (1 :1000, sc-47778, Santa Cruz), GAPDH (1:1000, AM4300, Invitrogen). After three washing steps of 10min with TBS-T, membranes were incubated with horseradish peroxidase (HRP)—conjugated anti-mouse or anti-rabbit secondary antibodies for 1 h at RT: Goat Anti-Mouse IgG HRP (1:4000, 10004302, Cayman Chemicals) or Goat Anti-Rabbit IgG HRP (1:10.000, 1706515, Bio-Rad). After three washing steps of 10min with TBS-T, chemiluminescent detection was performed with enhanced chemiluminescence (ECL) western blotting detection reagent (GE Healthcare). For AICD detection, longer exposure times were applied. Optical density was determined with Image-J, according to the software instructions (T. Ferreira. & Rasband, 2012). Full-length blots with the molecular weight standards (NZYColour Protein Marker I, NZYTech) are provided in **Supplementary Figures 5 and 6**.

### Primary neuronal cultures

Hippocampal neurons were cultured from 18-day Sprague-Dawley rat embryos, adapting the protocol from (Temido-Ferreira et al., 2020, Afonso et al., 2019). Briefly, embryos were collected in Hank’s Balanced Salt Solution (HBSS, Corning) and rapidly decapitated. After removing the meninges, hippocampi were dissociated in HBSS with 0.25% trypsin at 37°C for 15 min, resuspending every 3min. The tissue was then washed with HBSS containing 30% fetal bovine serum (FBS) to stop trypsin activity, followed by three washing steps with HBSS. Cells were resuspended in neuronal plating medium (MEM (Minimum Essential Medium) supplemented with 10% horse serum, 0.6% glucose, and 100 U/mL Pen-Strep), gently dissociated and filtered through a 70μm strainer. Finally, cells were plated on poly-D-lysine-coated glass coverslips (0.1mg/mL PDL in 0.1M borate buffer, pH 8.5) in 24-multi well plates at a final density of 70.000 cells/coverslip, in neuronal plating media and maintained at 37°C in a 5% CO_2_-humidified incubator. After 4 hours, the plating medium was replaced for neuronal culture medium: Neurobasal Medium (Gibco–Life Technologies) supplemented with B-27 supplement, 25μM Glutamic acid, 0.5mM Glutamine, and 20 U/ml penicillin/streptomycin. Cultures were maintained in the humidified incubator for 2 weeks, feeding the cells once per week with neuronal culture medium by replacing half of the medium per well.

### Neuronal transfection

Primary neuronal cultures were transiently transfected at DIV (days *in vitro*) 7-8 using the calcium phosphate transfection protocol adapted from (Silva et al., 2019, Jiang et al., 2004). For each well, 1.5µg plasmid DNA was diluted in 17.5µL TE buffer (10 mM Tris, 1 mM EDTA, pH 7.3). CaCl_2_ solution (2.5M in 10mM HEPES, pH 7.2) was added dropwise to the diluted DNA (final concentration = 250 mM CaCl_2_) and gently mixed. This mix was then added dropwise to an equivalent volume of HEPES-buffered saline transfection solution (in mM: 274 NaCl, 10 KCl, 1.4 Na_2_HPO_4_, 11 dextrose, 42 HEPES, pH 7.2), gently mixed and incubated at room temperature for 30 min, vortexing every 5 min. During this period, neurons were treated with 2 mM kynurenic acid in conditioned neuronal culture medium (in a new multi-well plate). The precipitates were then added dropwise to pre-conditioned neurons, followed by an incubation of 2-3h at 37°C in a 5% CO_2_-humidified incubator. Finally, DNA precipitates were dissolved by incubating the neurons in an acidified neuronal culture medium (in mM: 2 kynurenic acid, ∼5 HCl final concentration) for 15-20 min at 37°C. Coverslips were transferred to the original plates with conditioned neuronal culture media maintained in the humidified incubator.

### Plasmid generation

Primary neuronal cultures were transfected with AAV-shRNA–mCherry plasmids, with a shRNA against APP or a shRNA control sequence.

The control plasmid was kindly provided by Dirk Grimm (University of Heidelberg) and corresponds to AAV-U6-NS1-CMV-mCherry plasmid, where NS1 is a non-silencing sequence: GTAACGACGCGACGACGTAA, with no identified targets in the rat genome, confirmed by NCBI Basic Local Alignment Search Tool (BLAST).

For the generation of the shRNA-APP construct, we used the following sequence: GCACTAACTTGCACGACTATG (Young-Pearse et al., 2007), which is complementary to the mRNA NCBI reference sequence for rat APP (Rattus norvegicus amyloid beta precursor protein (App), NM_019288.2). This construct, which was provided by Tracy Young-Pearse (Harvard Medical School) in the pENTR-U6 vector, was then subcloned into an adeno-associated virus backbone (AAV-U6-shRNA empty-CMV-mCherry plasmid), kindly provided by Dirk Grimm. Briefly, the shRNA insert was generated by PCR amplification using primers with AscI and XhoI restriction sites (Forward Primer: AscI-U6 5’-GCGGCGCGCCAGGAAGAGGGCCTATTTCCCATG-3’; Reverse Primer: XhoI-PolyA-Active 5’-GCAAGTTAGTGCTTTTTTCTAGACCCTCGAGCG-3’). Subsequently, the PCR product and the AAV-U6-shRNAempty-CMV-mCherry plasmid were digested with AscI and XhoI restriction enzymes. Following gel purification, the shRNA construct was ligated into the AAV plasmid and the ligation product was transformed into Top10 chemically competent cells. Both plasmids were purified using the EndoFree Plasmid Maxi Kit (Qiagen) and verified by DNA sequencing (Primer 5’-GGGCCTATTTCCCATGATTCC-3’).

### Immunocytochemistry

The immunostaining protocol was adapted from (J. S. Ferreira et al., 2017). Briefly, neurons were fixed at DIV 14-15 (7 days after transfection) in 4% sucrose and 4% paraformaldehyde in phosphate-buffered saline (PBS) for 20 min at room temperature (RT). Neurons were then washed 3 times with PBS and permeabilized with PBS + 0.25% (v/v) Triton X-100 at RT (for 10min in the case of intracellular epitopes and 5 min for extracellular targets). Following 3 washing steps in PBS, cells were incubated in 10% (w/v) bovine serum albumin (BSA) in PBS for 1 h at RT to block nonspecific staining. Incubation with primary antibodies was performed overnight at 4°C in a humidified chamber, with antibodies diluted in 3% BSA PBS: APP Y188 (1:100, ab32136, Abcam), GluN2B (1:100, AGC-003, Alomone), GluN2A (1:100, AGC-002, Alomone), PSD-95 (1:50, ADI-VAM-PS002-E, Enzo). Following 4 washing steps in PBS, cells were incubated with the appropriate secondary antibody diluted in 3% BSA PBS (1:500) for 1h at RT: Donkey anti-Rabbit IgG Alexa Fluor 488, Donkey anti-Mouse IgG Alexa Fluor 488 or Donkey anti-Rabbit Alexa Fluor 647 (Thermo Fisher). Finally, cells were stained with Hoechst 33258 (12 ug/mL in PBS, Life Technologies) for 5min, washed 3 times with PBS and mounted in Fluoromount aqueous mounting medium.

### Microscopy imaging and analysis

All images were acquired in a Zeiss LSM 880 laser scanning confocal microscope using a Plan-Apochromat 63x/1.4 oil immersion objective.

For the analysis of APP immunofluorescence in transfected neurons, Hoechst fluorescence was detected using 405 nm for excitation (Diode laser with 30 mW nominal output – 2% transmission) and a 415-475 nm detection window, with PMT gain set to 610 and offset to −1. Alexa Fluor 488 fluorescence was detected using the 488 nm laser line of an Ar laser for excitation (25 mW nominal output – 1% transmission) and a 498-557 nm detection window, with GaAsP detector gain set to 500 and offset to 1. mCherry fluorescence was detected using 594 nm for excitation (HeNe laser with 2 mW nominal output – 5% transmission) and a 600-735 nm detection window, with PMT gain set to 700 and offset to 1. The pinhole size was set to 1.67 AU for Hoechst, 1.37 AU for Alexa Fluor 488 and 1.1 AU for mCherry. Z-stacks of the three channels were acquired with Zoom set to 1 (134.95×134.95 µm area with 1024×1024 pixel frame size - 0.13 µm pixel size) with a 0.49 µm slice interval, a line average of 2 and 1.03 µs pixel dwell time (unidirectional scan). The APP (Alexa Fluor 488) relative fluorescence intensity was manually quantified using ImageJ, after maximum intensity projection. For each condition, 7 transfected neurons were analyzed by defining regions of interest (ROI) which corresponded to the cell bodies using the mCherry channel. The average intensity of Alexa Fluor 488 was then determined for each ROI. All values were normalized to the average intensity in transfected neurons from the control condition (%).

For the analysis of GluN2B/PSD95 in dendrites of transfected neurons, Alexa Fluor 488 fluorescence was detected using the 488 nm laser line of an Ar laser for excitation (25 mW nominal output – 3% transmission) and a BP 495-550 + LP 570 nm filter set for detection in the Airyscan unit, with gain set to 790 and offset to 0. mCherry fluorescence was detected using 561 nm for excitation (DPSS laser with 20 mW nominal output – 7.5% transmission) and a BP 495-550 + LP 570 nm filter set in conjunction with a SP 615 nm secondary beam splitter for detection, with gain set to 850 and offset to 0 in the Airyscan unit. Alexa Fluor 647 fluorescence was detected using 633 nm for excitation (HeNe laser with 5 mW nominal output – 14.5% transmission) and a BP 570-620 + LP 645 nm filter set in conjunction with a LP 660 nm secondary beam splitter for detection with gain set to 760 and offset to 0 in the Airyscan unit. The pinhole size was set to 2.66 AU for Alexa Fluor 647, 2.16 AU for mCherry and 2.49 AU for Alexa Fluor 488. Z-stacks of the three channels were acquired with Zoom set to 3.8 (35.51×35.51 µm area with 836×836 pixel frame size - 0.04 µm pixel size) with a 0.18 µm slice interval, a line average of 2 and 2.53 µs pixel dwell time (unidirectional scan). All data sets were subjected to Airyscan processing in ZEN using the same parameters. The analysis of GluN2B/PSD95 in transfected neurons was performed using an in-house developed macro for ImageJ. For each condition, 13 dendrites from transfected neurons were analyzed. Following maximum intensity projection and manual selection of the dendritic area using the mCherry channel, images were segmented with user-defined intensity thresholds for GluN2B (Alexa Fluor 647) and PSD95 (Alexa Fluor 488), which were maintained constant for all conditions. The mean fluorescence intensity and the percentage of dendritic area with positive signal (above threshold) were quantified for each channel in the segmented images. GluN2B and PSD95 clusters were detected by particle analysis, which also allowed us to quantify average cluster sizes in each case. The colocalization area was determined by identifying the pixels where both GluN2B and PSD95 intensity values were above the respective threshold. The relative GluN2B synaptic content was then determined as the ratio between the area occupied by colocalized pixels and the total area with GluN2B staining. The fluorescence density was calculated as the mean fluorescence intensity multiplied by the percentage of dendritic area with positive signal for each channel. All quantifications were normalized to the average values of transfected dendrites from the control condition (%).

## Statistical Analysis

All statistical analyses were performed with GraphPad Prism software. Results are presented as dot blots with individual values plus bar, with mean ± Standard Error of the Mean (SEM). Statistical analyses were performed after evaluating normality using the Shapiro–Wilk test. When the distribution was normal in all groups, the statistical comparison included unpaired t test or one-way ANOVA followed by multiple comparison tests. Uncorrected Fisher’s LSD’s multiple comparisons test was used when comparing the mean of one group with the mean of another. Tukey’s multiple comparisons test was used when comparing the mean of more than 2 groups. When the distribution was not normal, the statistical comparison was performed by Mann-Whitney test or Kruskal Wallis followed by multiple comparisons tests. Uncorrected Dunn’s test was used when comparing the mean of one group with the mean of another. Correlations between parameters were determined according to Pearson’s correlation coefficient. Significance was determined according to the following criteria: p>0.05= not significant, *p<0.05, **p<0.01 ***p<0.001 and ****p< 0.0001. The complete statistical values are provided in **Supplementary Table 1**.

## Data availability

For detailed information on experimental design please see the provided Reproducibility Checklist. All the software used to data analysis is commercially available and the respective information is provided in each respective section. The data that support the findings of this study are available from the corresponding authors upon request.

## Funding and acknowledgements

JR-S is an FCT/PhD Fellow (IMM Lisbon BioMed PhD program; SFRH/BD/52228/2013); LVL is supported by Fundação para a Ciência e a Tecnologia (PTDC/BIM-MEC/47778/2014). LVL is an FCT-CEEC Principal Investigator. AR, LVL and PP were supported by Santa Casa da Misericórdia (MB-7-2018). SM and HM were supported by supported by the FLAG-ERA grant (MILEDI) by ANR to SM and HM. We would also like to thank Dirk Grimm and Kathleen Börner (University of Heidelberg) and Tracy Young-Pearse (Harvard Medical School) for providing the plasmids; and to the Rodent and Bioimaging Facilities of Instituto de Medicina Molecular João Lobo Antunes for their technical support and the Biobanco-IMM, Lisbon Academic Medical Center, Lisbon, Portugal.

## Author contributions

JR-S has written the draft and performed the neuronal cultures, shAPP plasmid cloning, Western blotting and the immunocytochemistry assays. PAP performed the whole cell patch-clamp recordings. MT-F and JEC prepared primary neuronal cultures. MW provided tools to perform western blot analysis of APP processing. HM designed and cloned shAPP plasmids. AR and SM performed the in-vivo pharmacological treatment of aged mice. JRS, LVL and PAP designed the experiments and wrote the manuscript. LVL and PAP coordinated the project. All authors revised the manuscript and discussed the experimental findings.

---

**Supplementary Figure 1–.**
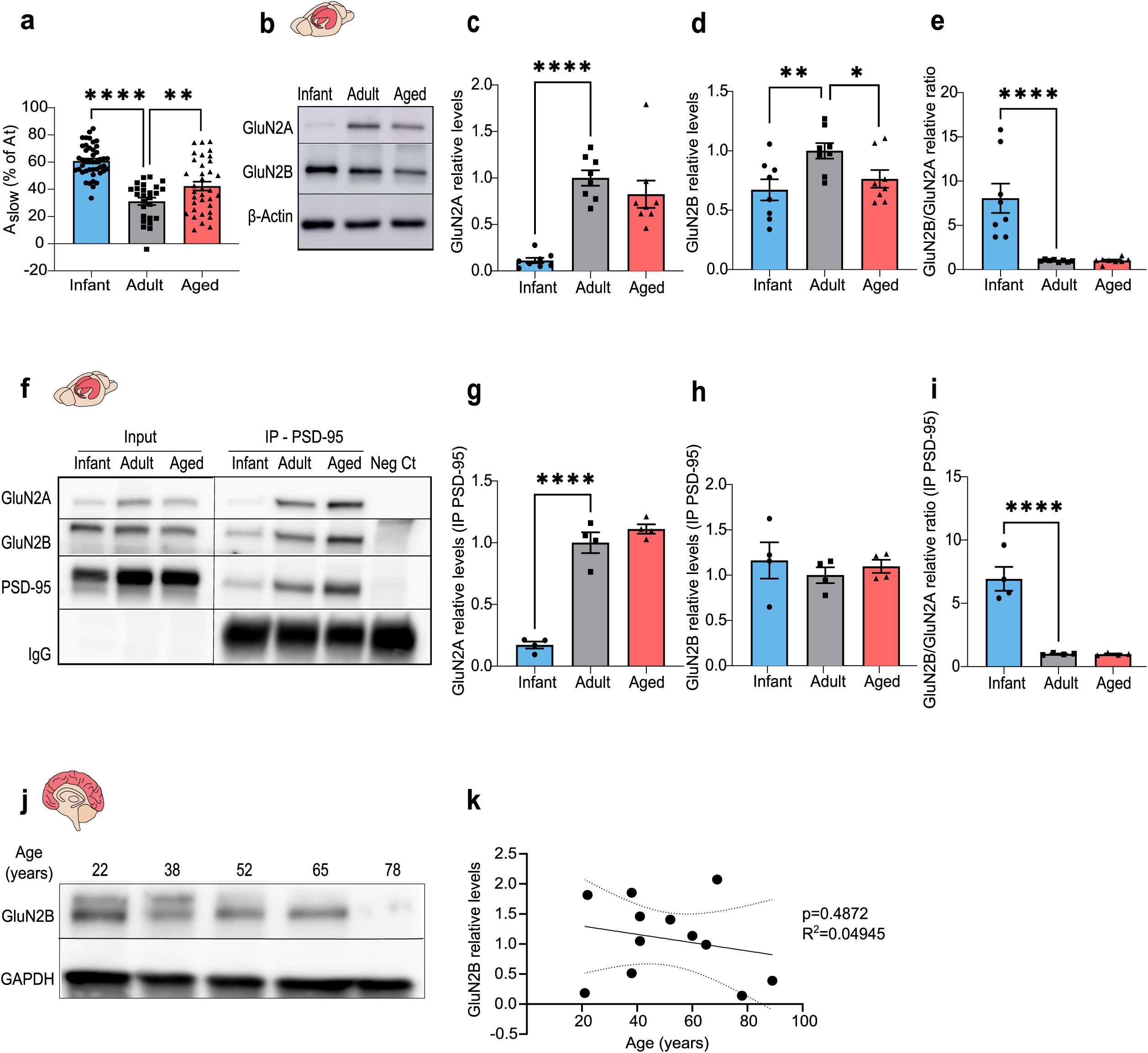
GluN2B/GluN2A ratio is increased in infant mice in hippocampal lysates and fractions immunoprecipitated for PSD95. (a) A_slow_ was calculated as the amplitude of the slow component of NMDAR EPSCs normalized to the total amplitude (%), measured by whole-cell patch-clamp recordings in CA1 hippocampal neurons from infant, adult and aged C57BL/6 mice. Results are expressed as the mean ± SEM (One-way ANOVA followed by an Uncorrected Fisher’s LSD’s multiple comparisons test using the adult group as reference, **p<0.01,****p<0.0001, n=28-44). (b) Representative western of hippocampal lysates from infant, adult and aged C57BL/6 wild-type mice. Membranes were immunoblotted with antibodies for GluN2A, GluN2B and β-actin. (c) Results from blots as shown in (b) from hippocampal lysates were normalized with β-actin and are expressed as the mean ± SEM (Kruskal Wallis test followed by Uncorrected Dunn’s test using the adult group as reference, ****p<0.0001, n=8). (d) Results from blots as shown in (b) from hippocampal lysates were normalized with β-actin and are expressed as the mean ± SEM (One-way ANOVA followed by an Uncorrected Fisher’s LSD’s multiple comparisons test using the adult group as reference, **p<0.01, *p<0.05, n=8). (e) Results from blots as shown in (b) from hippocampal lysates show the GluN2B/GluN2A relative ratio and are expressed as the mean ± SEM (One-way ANOVA followed by an Uncorrected Fisher’s LSD’s multiple comparisons test using the adult group as reference, ****p<0.0001, n=8). (f) Representative western blot of hippocampal lysates from infant, adult and aged C57BL/6 wild-type mice immunoprecipitated for PSD-95. Membranes were immunoblotted with antibodies for GluN2A, GluN2B and PSD-95. (g, h) Results from blots as shown in (f) from PSD-95 immunoprecipitated samples were normalized with PSD-95 and are expressed as the mean ± SEM (One-way ANOVA followed by an Uncorrected Fisher’s LSD’s multiple comparisons test using the adult group as reference, ****p<0.001, n=4). (i) Results from PSD-95 immunoprecipitated samples show the GluN2B/GluN2A relative ratio expressed as the mean ± SEM (One-way ANOVA followed by an Uncorrected Fisher’s LSD’s multiple comparisons test using the adult group as reference, ****p<0.0001, n=4). (j) Representative western blot of prefrontal cortex human samples (21 to 89 years old). Membranes were immunoblotted with antibodies for GluN2B and GAPDH. (k) Linear regression graph calculated from blots as shown in (j) shows the variation in GluN2B relative levels (normalized with GAPDH) depending on the age of human subjects (n=12). Statistical analysis was performed using Pearson’s correlation (two-tailed p value). Dotted lines represent the 95% confidence intervals. The values obtained for 20–25-year-old subjects were used as reference.

**Supplementary Figure 2.**
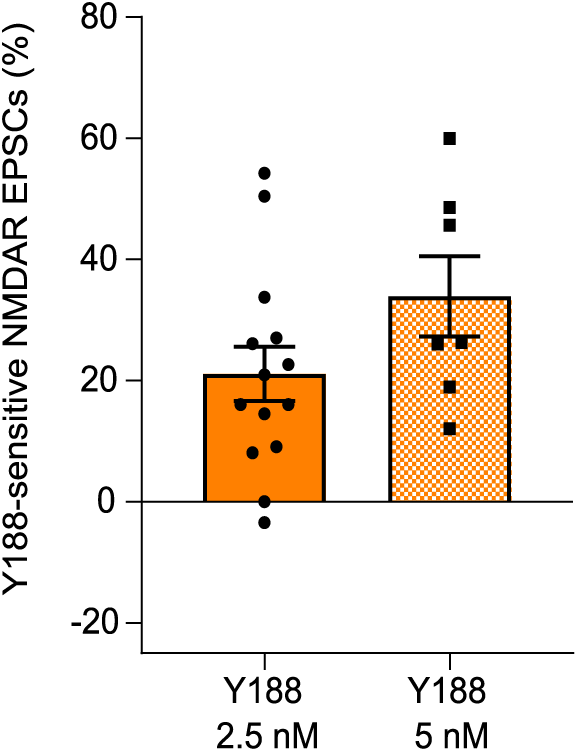
– Dose-response of APP C-terminal antibody in NMDAR EPSCs. The effect of the presence of the APP C-terminal antibody in the intracellular space in neurons from infant mice was evaluated using two concentrations of APP C-terminal antibody (Y188): 2.5 nM and 5 nM, introduced in the recording pipette. NMDAR EPSC amplitude was measured by whole-cell patch clamp in CA1 pyramidal neurons of infant C57BL/6 wild-type mice during 60 min of incubation with the antibody. The percentage of Y188-sensitive NMDAR EPSCs was calculated comparing the baseline amplitude (15-20min) with the final amplitude (60min) and normalized to the control condition (boiled Y188). Results are expressed as the mean ± SEM (Unpaired t-test, n=7-14).

**Supplementary Figure 3–.**
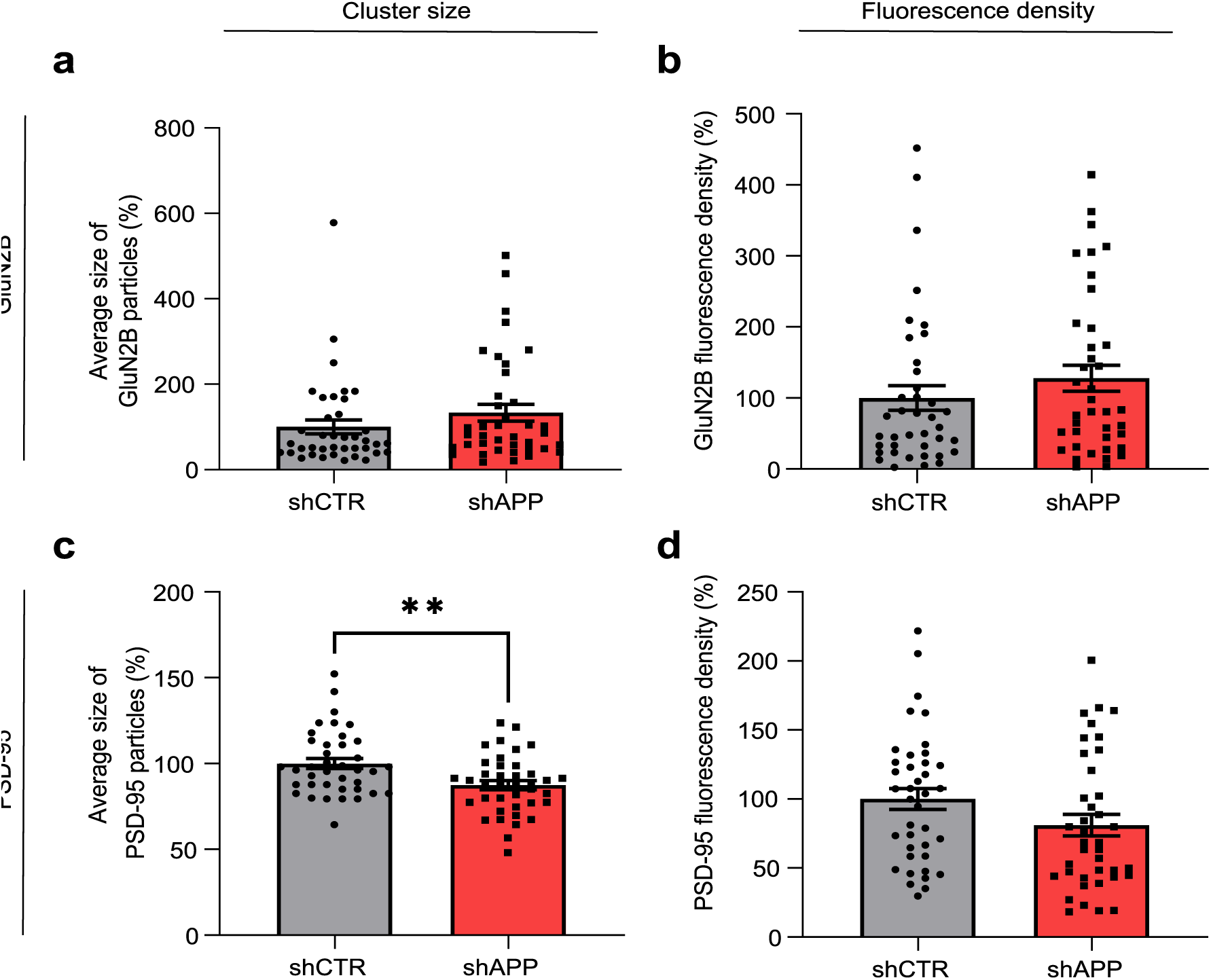
APP regulates PSD-95 but not GluN2B cluster size and has no impact on GluN2B/PSD-95 total levels. (a, b) Quantification of GluN2B average cluster size and fluorescence density in primary neuronal cultures (DIV14) transfected with shAPP or the respective control (shCTR) at DIV7. Results are expressed as the mean ± SEM, using the control condition as reference (%) (Mann-Whitney test, n=39 dendrites, 3 independent cultures). (c) Quantification of PSD-95 average cluster size in primary neuronal cultures transfected with shAPP or the respective control (shCTR). Results are expressed as the mean ± SEM, using the control condition as reference (%) (Unpaired t-test, n=39 dendrites, 3 independent cultures, **p<0.01). (d) Quantification of the relative PSD-95 fluorescence density in primary neuronal cultures transfected with shAPP or the respective control (shCTR). Results are expressed as the mean ± SEM, using the control condition as reference (%) (Mann-Whitney test, n=39 dendrites, 3 independent cultures).

**Supplementary Figure 4–.**
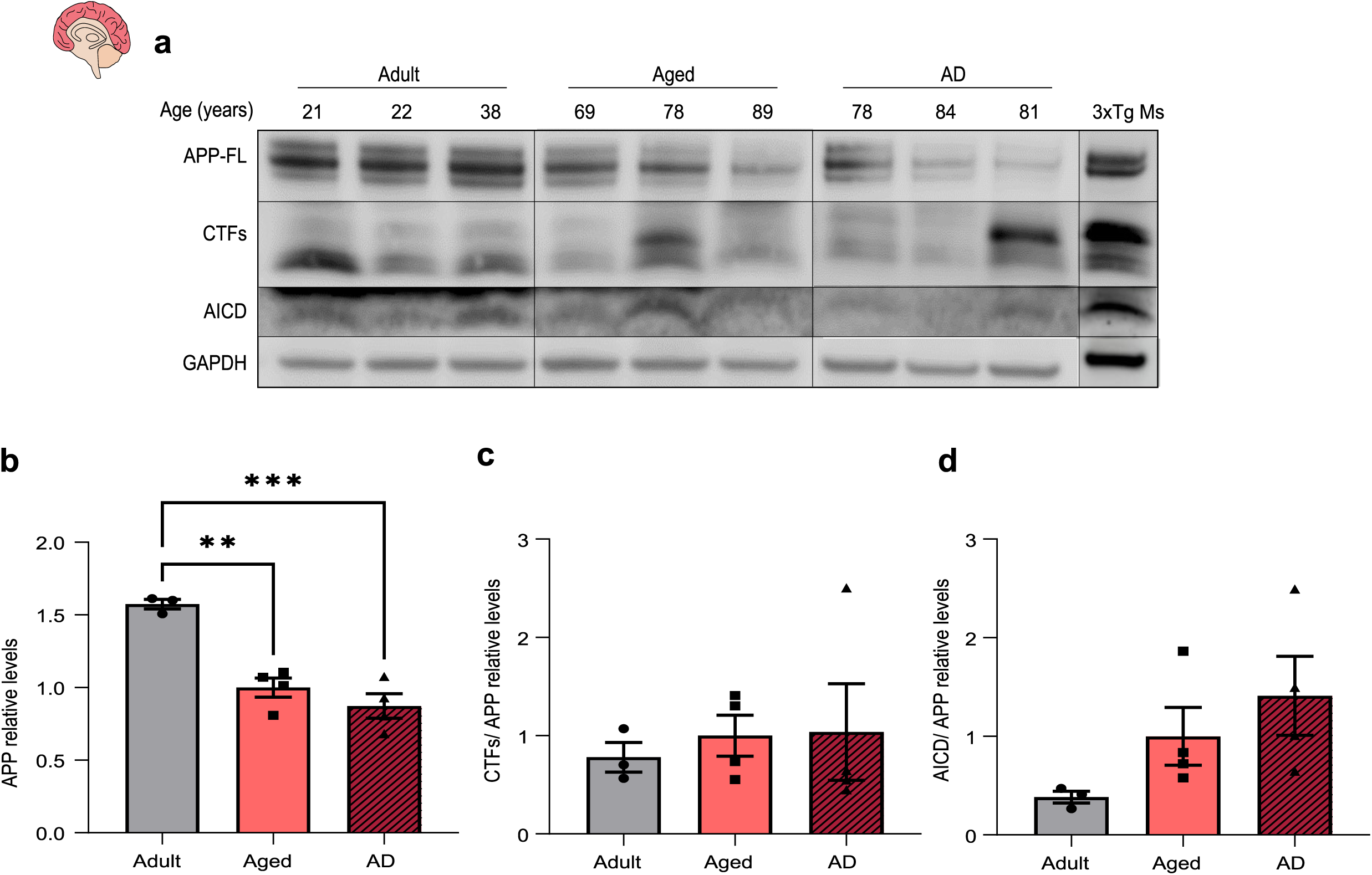
Alzheimer’s Disease patients exhibit a similar profile of APP levels/processing than aged controls. (a) Representative western blot of prefrontal cortex human samples (21 to 89 years old), with or without Alzheimer’s Disease (AD). Membranes were immunoblotted with antibodies for APP C-terminal (to detect APP full-length (APP-FL), APP C-terminal fragments (CTFs) and the APP Intracellular Domain (AICD)) and GAPDH. A female triple transgenic mouse (3xTg, 6 months) was used as a positive control for APP-derived fragments. (b) Quantification of APP relative levels in human samples (normalized to GAPDH) calculated from blots as shown in (a), comparing adult subjects (<40 years old), aged subjects (≥65 years old) and AD patients (≥65 years old). Results are expressed as the mean ± SEM, using the aged subjects as reference (One-way ANOVA followed by Tukey’s multiple comparisons test, ***p<0.001, **p<0.01, n= 3-4). (c) Quantification of CTFs/APP relative levels in human samples calculated from blots as shown in (a), comparing adult subjects (<40 years old), aged subjects (≥65 years old) and AD patients (≥65 years old). Results are expressed as the mean ± SEM, using the aged subjects as reference (Kruskal Wallis followed by Dunn’s multiple comparisons test, n=3-4). (d) Quantification of AICD/APP relative levels in human samples calculated from blots as shown in (a), comparing adult subjects (<40 years old), aged subjects (≥65 years old) and AD patients (≥65 years old). Results are expressed as the mean ± SEM, using the aged subjects as reference (One-way ANOVA followed by Tukey’s multiple comparisons test, n=3-4).

**Supplementary Figure 5 -.**
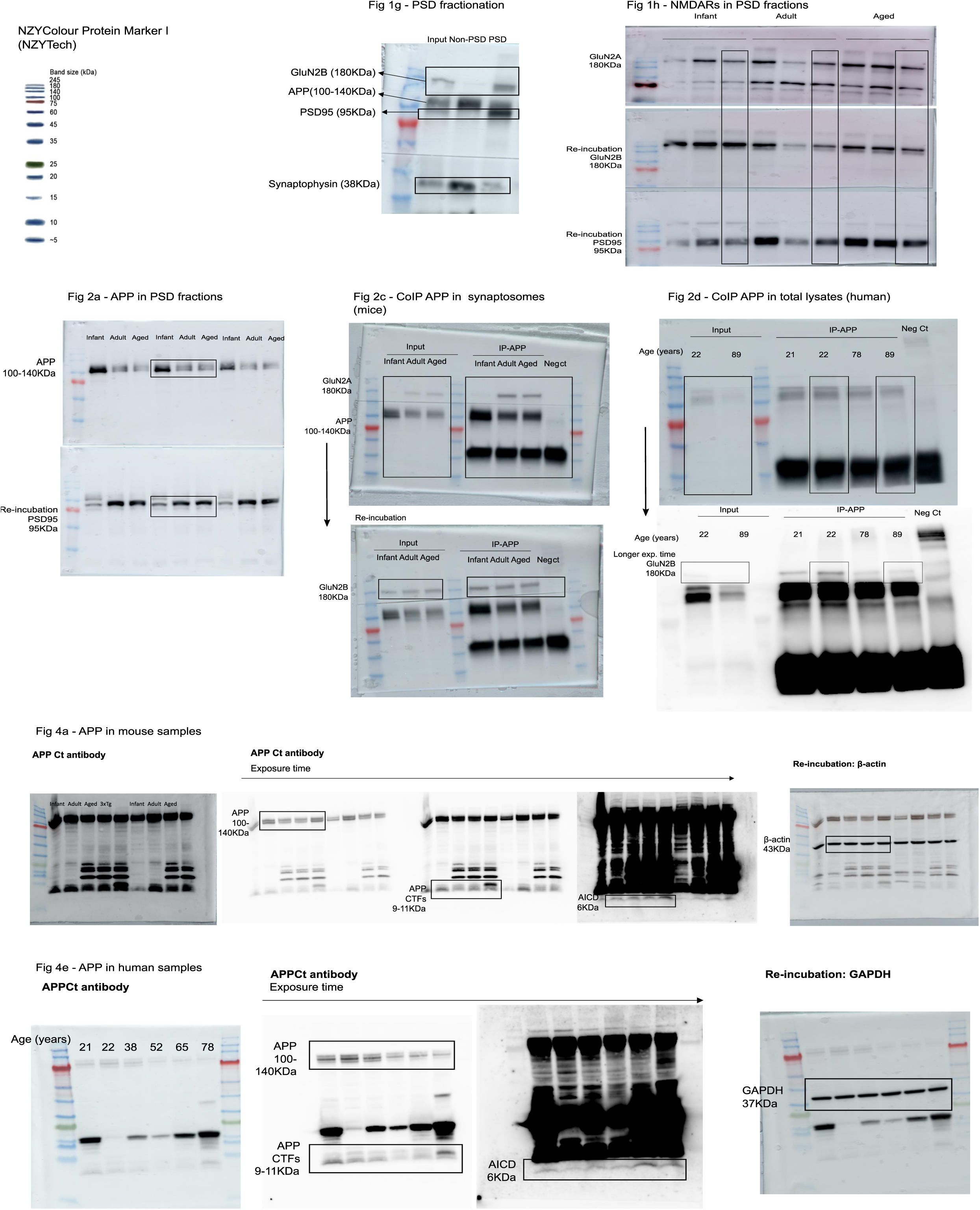
Full-length western blots with the molecular weight standards (NZYColour Protein Marker I, NZYTech) for the main figures.

**Supplementary Figure 6 -.**
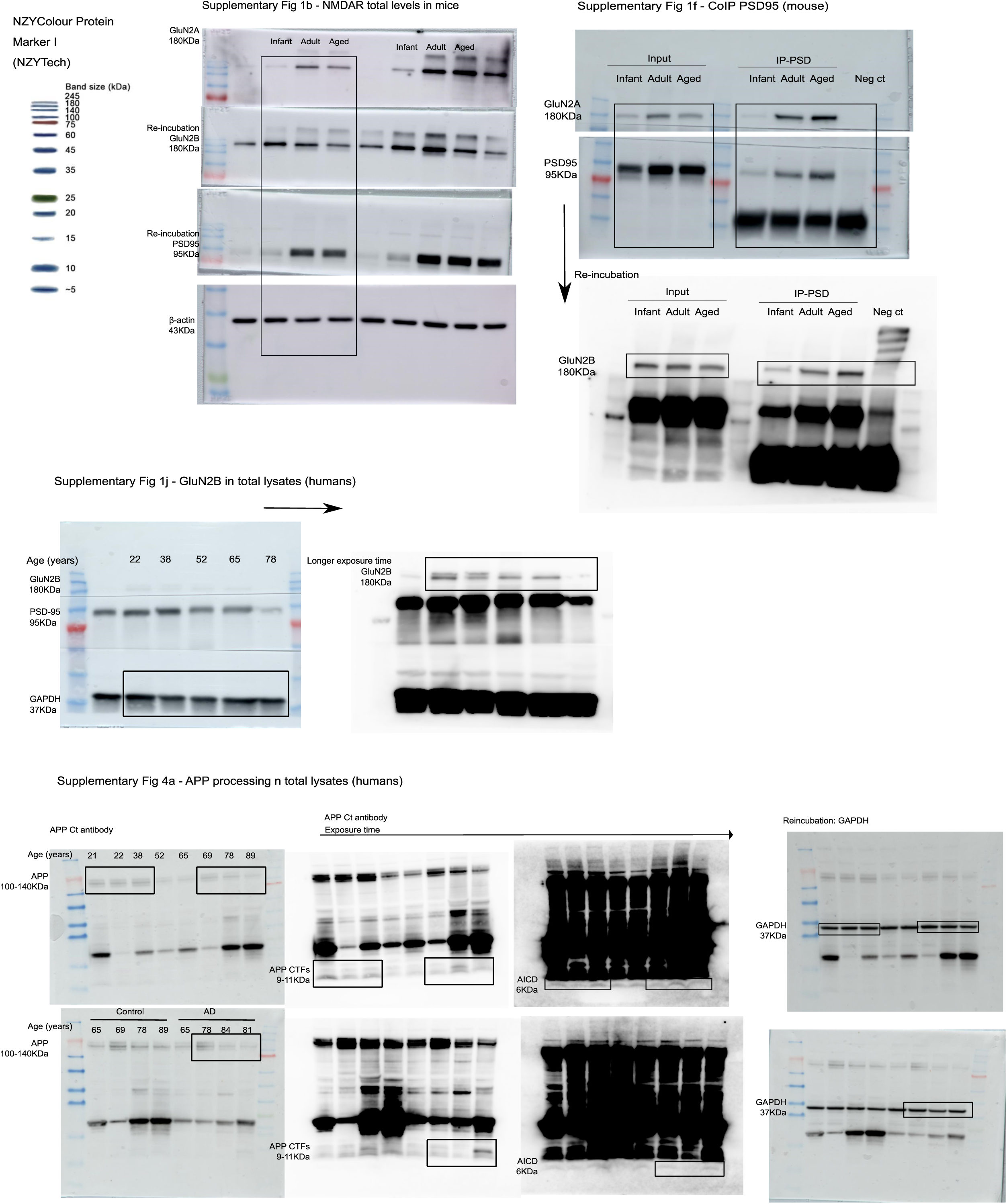
Full-length western blots with the molecular weight standards (NZYColour Protein Marker I, NZYTech) for the supplementary figures.

